# *DCX* knockout ferret reveals a neurogenic mechanism in cortical development

**DOI:** 10.1101/2024.06.13.598795

**Authors:** Wei Wang, Chonghai Yin, Shaonan Wen, Zeyuan Liu, Bosong Wang, Bo Zeng, Le Sun, Xin Zhou, Suijuan Zhong, Junjing Zhang, Wenji Ma, Qian Wu, Xiaoqun Wang

**Affiliations:** State Key Laboratory of Cognitive Neuroscience and Learning, IDG/McGovern Institute for Brain Research, Beijing Normal University, New Cornerstone Science Laboratory, Beijing, 100875, China; State Key Laboratory of Brain and Cognitive Science, Institute of Biophysics, Chinese Academy of Sciences, Beijing 100101, China; University of Chinese Academy of Sciences, Beijing 100049, China; Changping Laboratory, Beijing 102206, China; Beijing Institute of Brain Disorders, Capital Medical University, Beijing, 100069, China

**Keywords:** *DCX* knockout ferret, lissencephaly, snRNA-seq, neural progenitor cells, neurogenesis

## Abstract

Doublecortin (DCX) is one of the major causal proteins leading to lissencephaly and subcortical band heterotopia in human patients. However, our understanding of this disease, as well as the function of DCX during neurogenesis, remains limited due to the absence of suitable animal models that accurately represent human phenotypes. Here, we conducted a comprehensive examination of the neocortex at different stages in *DCX* knockout ferrets. We corroborated the neurogenic functions of *DCX* in progenitors. Loss of function of DCX led to the over-proliferation of neural progenitors and the truncation of basal processes of radial glial cells, which contributed to the thickening of cortices and the stalling of neurons underneath the cortical plate during neurogenic stages, respectively. We also present the first-ever cell atlas of the lissencephaly disease model, which embraces an almost reversed neuronal lamination distribution in the neocortex compared to the normal controls. Furthermore, we discovered alterations in molecular signatures tied to epilepsy, a condition frequently observed in lissencephaly patients. We also provided compelling evidence that the distribution of GABAergic inhibitory neurons in the cortex is intricately linked to glutamatergic excitatory neurons in a subtype-specific manner. In conclusion, our research offers new insights to expand our understanding of DCX’s functions and enrich our comprehension of lissencephaly’s intricacies.

**Highlights:** *DCX* ferrets phenocopy human lissencephaly and subcortical band heterotopia syndrome

*DCX* is required for NPC proliferation and radial glial basal fiber extension

The atlas of lissencephalic cortex is illustrated using snRNA-seq and spatial transcriptome

Inhibitory neurons couple to excitatory neurons in a cell-type specific manner

## INTRODUCTION

Lissencephaly (LIS) represents a range of gene-associated brain abnormalities characterized by a smooth cortical surface (due to the absence or reduction of gyri and sulci) in humans. This condition manifests through various symptoms, such as seizures, muscle spasms, developmental delays, cognitive impairments, and learning discrepancies, etc. ^1–3^ To date, nearly twenty gene mutations related to lissencephaly have been identified, with DCX being one of the most commonly reported risk genes in humans.^1–5^

The DCX gene, responsible for encoding Doublecortin, is located at Xq22.3 in humans. Hemizygous *DCX* mutations in males cause classical lissencephaly, while heterozygous mutations in females lead to subcortical band heterotopia (SBH).^6–10^ To illuminate the function of DCX in brains, great progress has been achieved by using animal models, *in vitro* cell culture, and induced pluripotent stem cells (iPSCs) derived from patients.^3,5,7,11–15^ *In vitro* studies identify that DCX likely directs neuronal migration by regulating the organization and stability of microtubules.^7^ However, it is surprising that *Dcx* knockout mice have shown no defect in cortical neuronal lamination or positioning.^13^ Although shRNA-mediated knockdown of Dcx in rats shows neuronal migration defects in the cortex,^12^ the specificity of the technique was challenged.^16^ However, other lissencephaly-related genes, such as *Reln*^17,18^ and *Lis1* (*Pafah1b1*)^19^, display severe cortical lamination defects in mice, which implies that DCX might work differently between humans and rodents.

Therefore, a new animal model that can faithfully replicate the phenotypes in human patients is urgently needed to further our understanding of the function of DCX in neural development as well as the profiling of lissencephaly.

Given the distinct and bewildering phenotypes between mice and humans, we propose that DCX might function differently in evolution. To address the function of DCX, and to better understand the cellular alterations and underlying mechanisms in lissencephaly brains, we generated *DCX* knockout ferrets with CRISPR/CAS9 technology.^20^ According to our de novo DNA sequencing and chromosome assembly, DCX is also located on the X-chromosome in ferrets, just as it is in humans (unpublished data). After several years of backcrossing, we successfully established two stable *DCX* knockout lines, both inducing a stop codon in the second exon (Figure S1A). The two lines exhibited the same phenotypes. Heterozygous females were fertile and viable; however, nearly all homozygous females died before adulthood. Only a few males survived to adulthood, and their fertility was very low. In this study, we used Line1 for further analysis. *DCX* knockout ferrets effectively reproduced the morphological and pathological characteristics observed in human patients, including classical lissencephaly and subcortical band heterotopia. We demonstrated unreported neurogenic functions of *DCX* in progenitors. We also observed that DCX is vital for the proper extension of radial glial fiber, which acts as a scaffold, guiding neurons to their terminal positions. In adulthood, the results of snRNA-seq, spatial transcriptomics, and immunofluorescence revealed the characteristics of the nearly ‘reversed’ cortex in DCX knockout ferrets. Additionally, we also found evidence indicating that inhibitory neurons preferentially are associated with specific excitatory counterparts during their migration and distribution in the neocortex, a process that was also impaired in *DCX* knockout ferrets.

## Results

### DCX regulates the proliferation of neural progenitor cells in ferret

Firstly, we confirmed the knockout animals using a specific antibody against DCX in both wild type (WT) and mutant ferrets (*DCX^-/-^*) at P7 (Figure S1B; Table S1). The results clearly showed an abundant DCX-positive signal in the cortical plate and the VZ and SVZ in WT, but no DCX-positive signal was detected in *DCX* mutant ferrets, indicating the successful knockout strategy. Interestingly, no DCX signal was detected in the VZ in mice at both E14.5 and E16.5 (Figure S1B), but highly expressed in neural progenitor cells in ferret brains (Figures S1B, S1C). DCX was also detected in the neural progenitors in developing macaque (E75) and human (GW18) brains (Figure S1D), which implies that DCX might have divergent functions among animals,^21^ particularly in NPCs.

Unlike primate brains, ferrets exhibit abundant cortical neurogenesis that persists into the postnatal stage, with cortical folding commencing from P4.^22^ Thus, we investigated the phenotypes of WT and *DCX* mutant (*DCX^-/-^*, *DCX^-/Y^*) at P0, a stage equivalent to E15.5-E16.5 in mice. To determine whether DCX affected neurogenesis, we used antibodies against neural progenitor markers, such as SOX2, HOPX, and TBR2, to delineate the expression profiles of RG, outer radial glial cell (oRG), and IPC in the somatosensory cortices, respectively (Figures 1A, 1B). HOPX is an oRG marker but is also detected in RGs in the VZ. For the quantification of HOPX^+^ cells, only cells in the oSVZ were counted. The results revealed an increase in SOX2^+^ progenitors in *DCX* knockout mutants compared to WT at P0. The number of IPCs (TBR2^+^) and oRGs (HOPX^+^) also significantly increased (Figures 1A-1C). We then examined the proliferation capacity of neural progenitors using Ki67 and pH3 (Phospho-Histone-H3). We observed more proliferating progenitors in *DCX* knockout brain cortices (Figures 1D, 1E, S1E, and S1F). At P7, we also observed more NPCs, proliferating NPCs, and the ratio of proliferating cells to NPCs (SOX2 and pH3 double positive, TBR2 and pH3 double positive, SOX2 and Ki67 double positive) in *DCX* knockout mutants, suggesting a more active neurogenesis (Figures 1F-1I, and S1G, S1H). In conclusion, we demonstrated that the loss of *DCX* increased the proliferation of neural progenitors, including RG, oRG, and IPC (Figure 1J).

**Figure 1.**
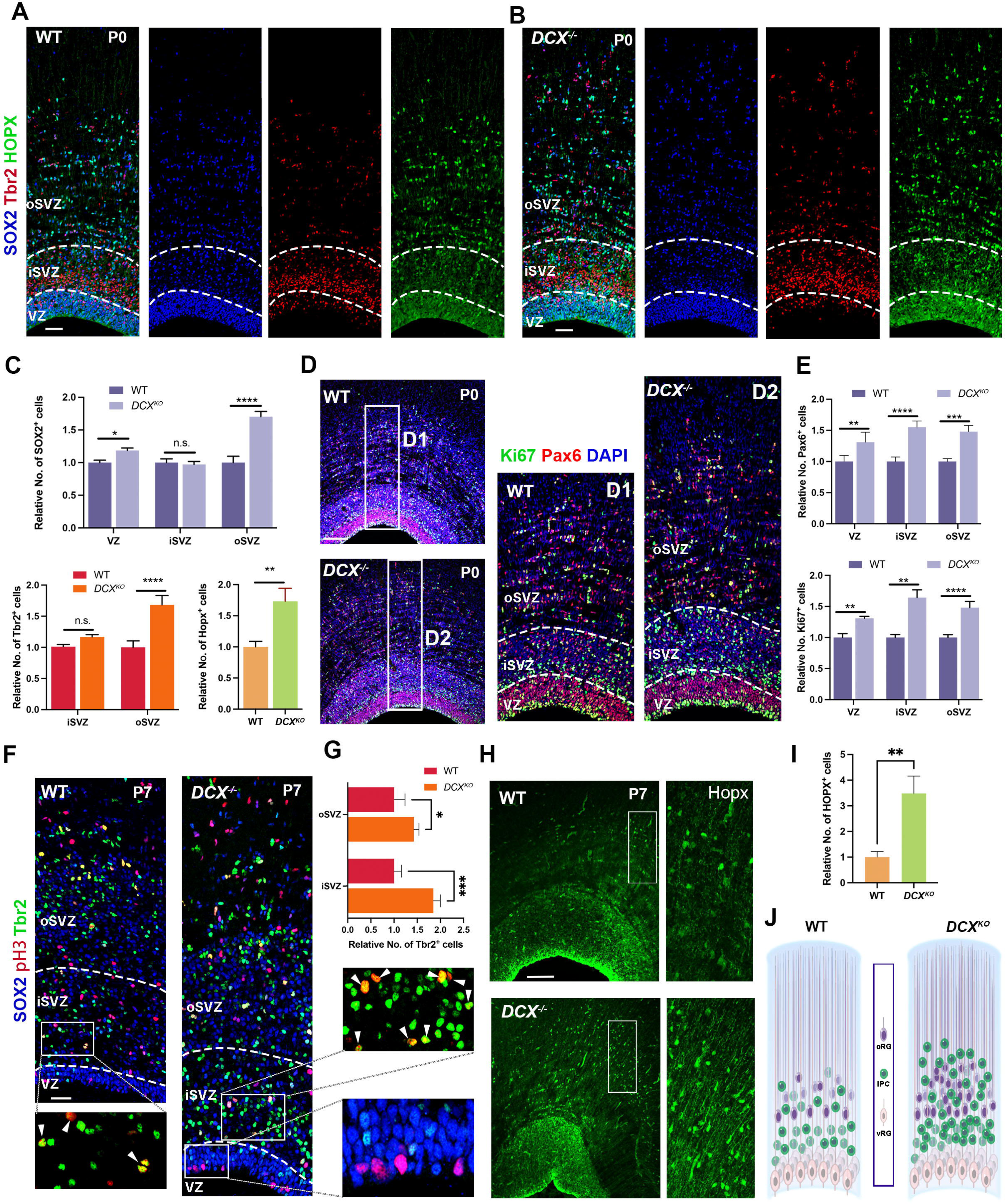
DCX suppressed the proliferation of neural progenitors. (A and B) The distribution of RG, oRG, and IPC viewed with SOX2, HOPX, and Tbr2 in the cortices of WT (n=5) and *DCX* (n=4) at P0. The VZ/SVZ/oSVZ was determined based on the expression of markers, cell density, and cell morphologies. (C) Quantification results showing the number of RG, oRG, and IPC increased in *DCX* knockout ferrets. (D and E) The number of proliferating cells is increased in *DCX* knockout ferrets. (n=4) compared to WT (n=5) at P0. (F-I) The number of neural progenitors (RG/oRG/IPC) and proliferating progenitors is increased in *DCX* knockout ferrets (n=3) compared to WT (n=3) at P7. (J) Schematic model showing the increased number of progenitors in *DCX* knockout ferrets. *p<0.05, **p<0.01, ***p<0.001, ****p<0.00001, unpaired two-tailed Student’s t-tests. Error bars represent SD. Scale bars, 100µm.

Next, we surveyed the mechanisms of DCX in regulating neural progenitor proliferation. We found that the cleavage plane orientation of mitotic RGs in the VZ shifted from vertical to horizontal (Figures 2A and 2B), which reportedly gives rise to proliferating progenitors and oRGs.^23,24^ The expression pattern of DCX in the VZ (Figure S1C) hinted that it might be involved in the regulation of radial glial cell junction and/or polarity. Then, we screened factors involved in adhesion junctions, tight junctions, and cell polarity complexes to identify potential factors that could interact with DCX to regulate the orientation of the cleavage plane. We observed a significant decrease in the expression of ZO-1 in DCX knockout mutant brains (Figures 2C and 2D), indicating a disruption in the polarity of RGs in DCX mutant animals. Therefore, the results suggested that the loss of DCX affected the proliferation of neural progenitors by disrupting the polarity and spindle orientation of RGs.

**Figure 2.**
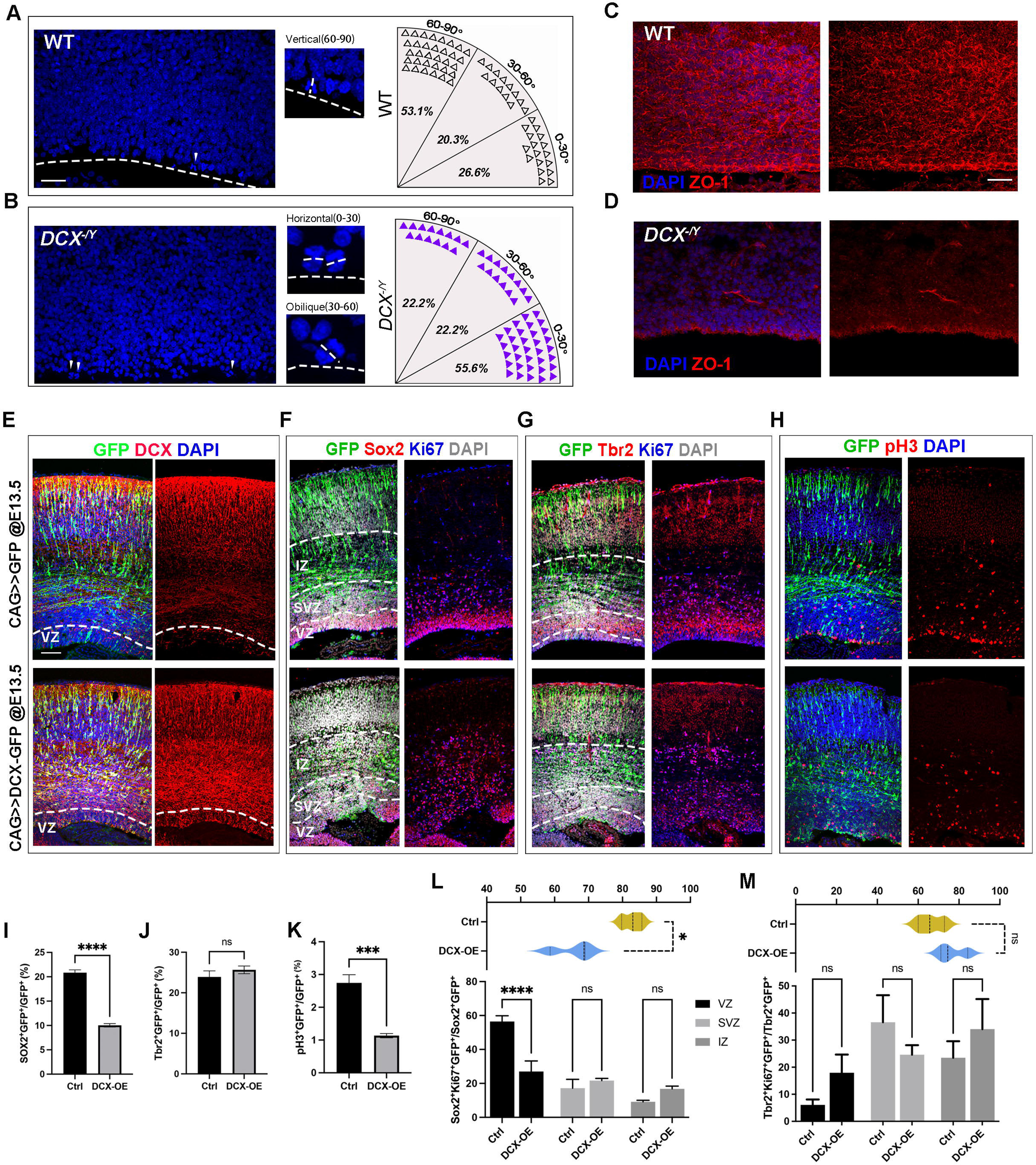
Knockout of DCX in ferrets and overexpression of DCX in mice validated the role of DCX in proliferation. (A-B) The division cleavage angle to the VZ surface in mitotic RG switched from vertical in WT to horizontal in *DCX* mutants. Scale bars, 100µm. (C-D) The expression of the cell junction and polarity protein ZO-1 is markedly decreased in *DCX* mutants compared to that in WT. Scale bars, 100µm. (E) Ectopic expression of DCX in the mouse neocortices by IUE. Scale bars, 100µm. (F-H) Ectopic expression of DCX in mice led to disorganized cortices and decreased the number and proliferation of apical progenitors. IZ: intermediate zone. (I-M) Quantification results of overexpression of DCX in the mouse cortices, compared to WT. *p<0.05, **p<0.01, ***p<0.001, ****p<0.00001, unpaired two-tailed Student’s t-tests. Error bars represent SD.

To confirm that DCX is essential for neural progenitor proliferation, we ectopically over-expressed DCX in the mouse neocortex using in utero electroporation (IUE). Ectopic DCX (pCAG-DCX-GFP) was introduced at E13.5, and the samples were collected and analyzed at E16.5. The results showed that DCX was ectopically expressed in the radial glial cells of mice (Figure 2E). We detected a decrease in the number of apical progenitors (Sox2^+^) in the cells with ectopic DCX expression (DCX-OE for short) in mice brains, but the number of Tbr2^+^ IPCs did not change significantly (Figures 2F, 2G, 2I, and 2J). Both Sox2 and Tbr2-positive cells exhibited disorganized patterns in DCX-OE. We observed that the number of pH3^+^ mitotic cells was dramatically decreased in DCX-OE compared to negative control (Figures 2H, 2K). A marked decrease in the ratio of proliferating apical progenitors in DCX-OE compared to WT (65.44% vs 82.84%, p = 0.0102) was also detected. When different regions were taken into account, cells in the VZ showed the most significant difference (Figure 2L). No significant difference was observed in the ratio of proliferating basal progenitors between DCX-OE and WT (76.64% vs 66.2%, p = 0.1273; Figure 2M). The proliferation rate of apical and basal progenitors is much higher than those in ferrets (∼40% and ∼30%, respectively; Figure S1F). The results highly support the function of DCX in modulating neural progenitor proliferation. In all, we concluded that overexpression of DCX in mice brains induced the delamination of progenitors and decreased the proliferation of neural stem cells, especially the RGs.

### Radial glial process defects contributed to the aberrant distribution of excitatory neurons in *DCX* knockout animals

The brain surface of ferrets is almost smooth at P0 and gradually develops gyri and sulci from P4.^22,25,26^ To investigate when the differences occured between WT and *DCX* knockout ferrets, we examined the brain cortices at P0 and P7. At P0, we observed the morphology and thickness of WT and *DCX* cortices were comparable, and the neuronal distribution appeared normal in *DCX* knockout brains (Figure 3A). Early-born neurons (TBR1/CTIP2-positive cells) were positioned in the deep region of the cortical plate, while a fraction of SATB2^+^ cells had migrated to the outermost region of the cortex, indicating that neuronal migration was not strongly affected at this stage. An elevated number of SATB2^+^ neurons was observed in the intermediate zone of *DCX* knockout ferrets (Figures 3A, 3B, and 1J). By P7, WT brains began to fold and expand laterally, while the mutant brain surface remained smooth (Figures 3C and S2A-S2C). Concurrently, the expression of layer-specific neuronal markers exhibited significant differences (Figures 3D-3F). There was an abundance of SATB2 and CUX1 positive cells stalled in the intermediate zone of *DCX* knockout cortices, compared to WT. This suggests an overproduction and migration defect of late-born upper layer neurons between P0 and P7, a critical time window that leads to the phenotypes observed in *DCX* knockout ferrets (Figures 3D-3F, S2A, and S2D).

**Figure 3.**
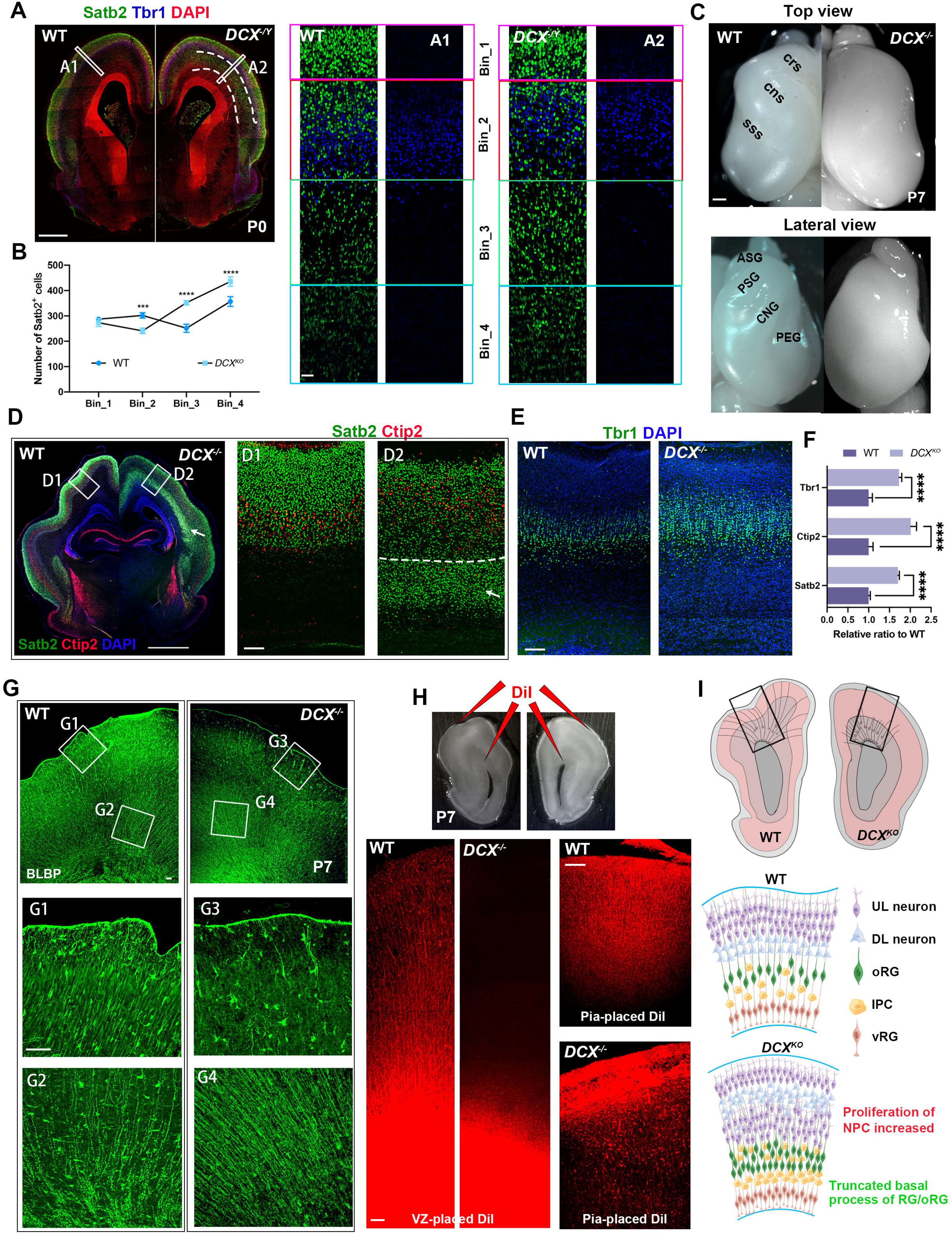
DCX is essential for the proper neuronal migration and extension of RG/oRG basal fibers. (A) Images of coronal hemibrain sections showing the expression patterns of SATB2^+^neurons and TBR1^+^ neurons in WT and *DCX* knockout ferrets at P0, dotted line indicates the subcortical band of SATB2^+^ neurons (scale bar, 1mm). A1 and A2 are divided into Bin 1 to Bin 4 (scale bar, 100µm). (B) Quantification results of A1 and A2 shows an increased number of SATB2^+^ neuron in Bin 3 and Bin 4 in *DCX* knockout ferrets. (C) Top and lateral views of brains in WT and *DCX* knockout ferrets at P7 (scale bar,1mm). WT brains show notable gyri and sulci on the surface. *DCX* knockout brains exhibit a lissencephalic phenotype. ASG, anterior sigmoid gyrus; CNG, coronal gyrus; cns, coronal sulcus; crs, cruciate sulcus; PEG, posterior ectosylvian gyrus; PSG, posterior sigmoid gyrus; sss, suprasylvian sulcus (D) Distribution of SATB2^+^ neurons and CTIP2^+^ neurons in WT and *DCX* knockout ferrets at P7 (scale bar, 2mm). The white arrow shows a subcortical band of STAB2^+^ neurons in *DCX*. Compared to WT (D1), an overwhelming accumulation of SATB2^+^ neurons were detected under the CTIP2^+^ neurons in *DCX* knockout ferrets (D2), suggesting a migration defect of SATB2^+^ neurons (scale bar, 100µm). (E) Images showing the number and laminar position of TBR1^+^ neurons in WT and *DCX* knockout ferrets (scale bar, 100µm). (F) Quantification results show that the number of SATB2^+^, CTIP2^+^, TBR1^+^ neurons is increased in *DCX* knockout ferrets compared to those in WT. (G) BLBP immunofluorescent staining in WT and *DCX* knockout ferrets at P7 (scale bar, 100µm). Weak BLBP-positive signals in *DCX* knockout ferrets (G3 and G4) are detected near the pial surface, compared to WT (G1 and G2). (H) DiI labeling assay showing the disintegrated basal processes of RG/oRG (scale bar, 100µm). Red needles indicate the placement locations of DiI crystals. (I) Schematic model showing DCX modulates the extension of RG/oRG fibers and the proliferation of NPCs.

During neurogenesis, newborn neurons migrate along the radial glial fibers to their appropriate destinations, a process orchestrated both intrinsically and extrinsically.^27,28^ DCX has been reported to play a role in neuronal migration through the regulation of microtubule organization and stability in neurons,^6,7^ which represents an intrinsic mechanism of neuron migration. The structural integration of radial glial fibers provides some of the extrinsic factors for neuronal migration. Thus, we investigated the architecture and organization of the radial glial basal fibers in both WT and *DCX* mutants. Immunostaining with Brain Lipid Binding Protein (BLBP) and Vimentin (VIM) revealed a well-organized radial glial fiber spanning from the ventral to the pial surface in both WT and *DCX* knockout brains at P0 (Figures S2E and S2F). However, at P7 we observed a significantly weaker or absent BLBP signal near the pial surface in *DCX* mutants, suggesting a discontinuity in basal fibers (Figure 3G, comparing G3 to G1). An immunofluorescence assay using VIM showed similar results (Figure S2G).

To confirm that the weakened or absent expression of both BLBP and VIM was a result of the disintegration of the basal processes of RG and oRG, DiI crystals were placed either in the VZ or at the pial surface (Figure 3H). In WT samples, DiI traced the path along the radial glial fibers from both sides, while in mutant samples, the DiI signal remained in the initial placement regions. This observation confirmed the discontinuity of the radial glial basal process in *DCX* knockout animals (Figure 3H). Taken together, these results demonstrate that lamination disorganization appeared at P7 but not at P0 in the developing cortex of *DCX* knockout ferrets. In terms of the underlying mechanism, our findings provide evidence that DCX plays a role in modulating the extension and maintenance of RG/oRG fibers, which act as scaffolds for migrating neurons. Thus, in addition to the well-established function of DCX in neurons, it also plays critical roles in neural progenitors, including the proliferation of NPCs and the extension of basal fibers in RG/oRG (Figure 3I). These results extended and broadened the function of DCX, which also provided a strong explanation for the undetectable defects in cortical lamination or positioning observed in mice.

### *DCX* Knockout Ferrets Replicate Human Phenotypes

In adults, WT ferret brains exhibited well-organized gyri and sulci (Figure 4A). *DCX* mutant brains displayed gender-specific phenotypes, similar to those reported in human patients,^6,9^ but with normal brain length and width compared to WT brains (Figure 4A). Male *DCX* mutants (*DCX^-/Y^*) exhibited a smooth brain surface without any folding (n=5, Figures 4A and 4B). In female *DCX* mutants (*DCX^+/-^*), we observed a subcortical band heterotopia phenotype (SBH) and a partial loss of sulci (n=3, Figures 4A and 4B). Coronal sections and Nissl staining images revealed that the parenchyma of brain cortices in *DCX^-/Y^* were cytoarchitecturally divided into an upper band and a deep band by a stripe of white matter (Figure 4B), forming a four-layered structure. The cortical thickness of mutant brains was also increased, mirroring the condition observed in human patients. Immunohistochemistry examination revealed a significant presence of NeuN^+^ cells in both the upper and deep bands in *DCX^-/Y^*and the SBH in *DCX^+/-^* samples, indicating the neuronal nature of these cells (Figure 4C). MRI imaging also verified that the brains of DCX mutants exhibit a loss folding on the surface throughout the entire brain (Figure 4D). The gyrification index ( GI ) score is 1.02 in *DCX^-/Y^* ferrets, compared to 1.467 in WT. In conclusion, our CRISPR/CAS9-mediated *DCX* knockout ferrets successfully recapitulated the lissencephaly or SBH phenotypes observed in humans, which provides an excellent disease model for studying the cellular and molecular alterations associated with these conditions (Figure 4E).

**Figure 4.**
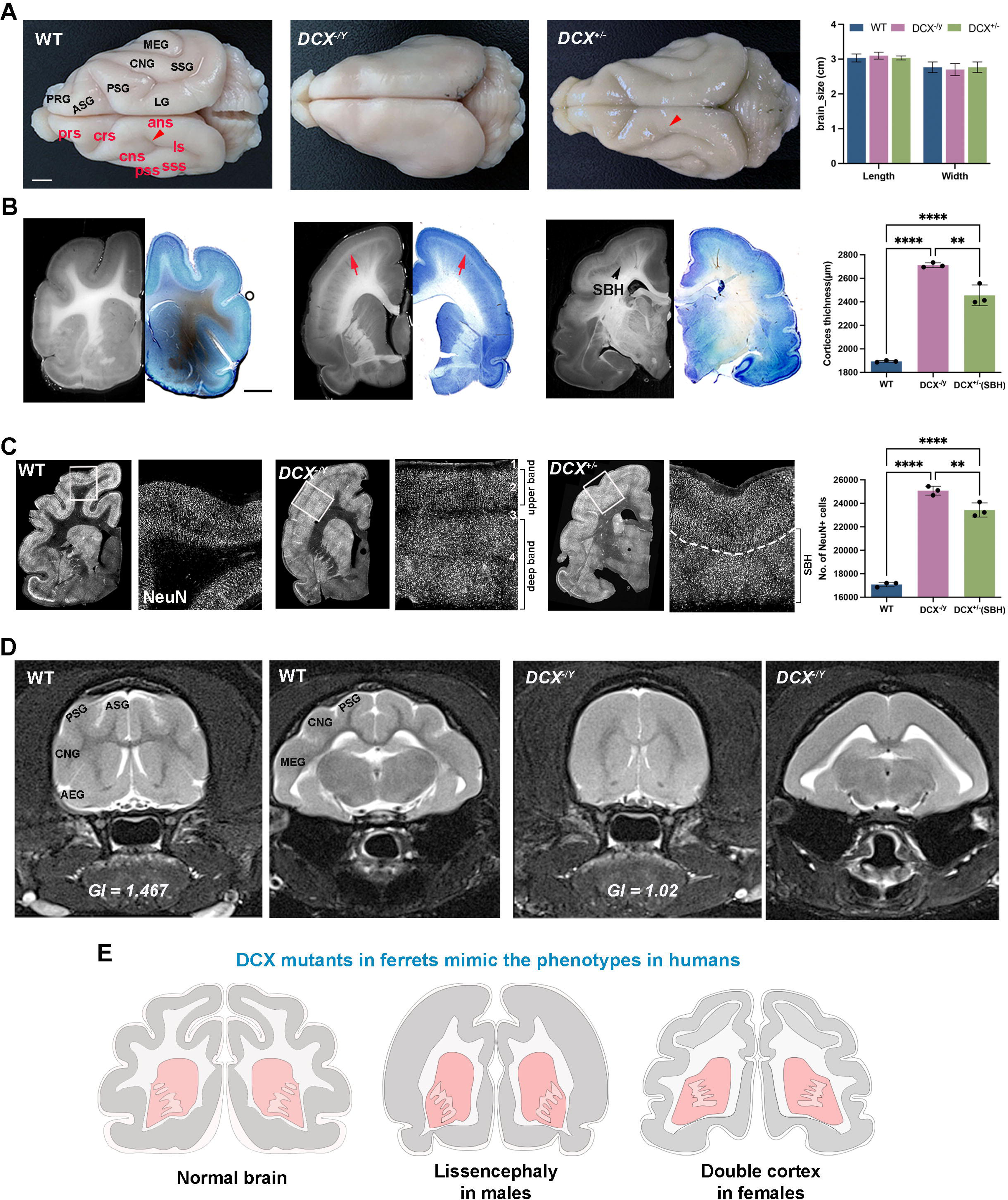
*DCX* knockout ferrets phenocopy human lissencephaly and SBH syndrome. (A) The phenotypes of *DCX^+/+^*, *DCX^-/Y^*, and *DCX^+/-^* ferret brains (n=3, scale bar, 3mm). WT brains exhibit exquisite gyri and sulci on the surface. *DCX* mutants show a typical lissencephalic phenotype in hemizygous males, and a mild loss of sulcus (ans, red arrow) in heterozygous females. ASG, anterior sigmoid gyrus; ans, ansate sulcus; CNG, coronal gyrus; cns, coronal sulcus; crs, cruciate sulcus; LG, lateral gyrus; ls, lateral sulcus; MEG, medial ectosylvian gyrus; PRG, proreal gyrus; PSG, posterior sigmoid gyrus; prs, prsylvian sulcus; pss, pseudosylvian sulcus; SSG, suprasylvian gyrus; sss, suprasylvian sulcus. (B) The morphology of coronal brain sections was observed using light microscope and Nissl staining, showing the phenotype of lissencephaly in hemizygous males and subcortical band heterotopia (SBH) in heterozygous females. The red arrow shows a stripe of white matter that separated the cortex into upper-band/deep-band in males; the black arrowhead indicates the SBH in females (n=3 in each group, scale bar, 500µm). (C) NeuN immunostaining results illustrate the neuronal identity and lamination of WT and *DCX* knockout cortices. The cortex was subdivided into a 4-layered structure instead of the 6-layered structure in *DCX* knockout ferrets (n=3 in each group). (D) MRI images showing lissencephaly in *DCX* ferrets. The GI scores are 1.467 and 1.02 in WT and *DCX* knockout ferrets, respectively. TR = 6580 ms, TE = 85 ms. (E) Schematic model of *DCX* knockout male and female ferrets. *p < 0.05, **p<0.01, ***p<0.001, ****p<0.00001, multiple comparison test after ANOVA. Error bars represent SD.

### Cellular landscape in WT and lissencephaly ferrets

To characterize the comprehensive molecular and cellular alterations in lissencephaly brains compared to WT brains, we dissected the cortices from controls, as well as the upper and deep bands in *DCX* male mutants (abbreviated as MU and MD, respectively, with n=2 biological individuals for each group) for snRNA-seq analysis (Figure 5A; Table S1). In total, we obtained 77,713 nuclei after stringent quality control and doublet removal (Figures S3A and S3B). Unsupervised clustering analysis and uniform manifold approximation and projection (UMAP) visualization were performed, grouping all cells into 24 clusters (Figure 5B). Based on the differentially expressed genes (DEGs) and classical markers, these 24 clusters were grouped into seven major cell types: glutamatergic neurons (ExNeu), GABAergic neurons (InNeu), oligodendrocyte precursor cells (OPC), oligodendrocytes (Olig), astrocytes (Astro), microglia (MG), and a combination of endothelial cells and pericytes (Endo_Peri) (Figures 5B, 5C, and S3C; Table S2). Additionally, cortical projection neurons (ExNeu_CPN) and callosal projection neurons (ExNeu_CFuPN) were identified. InNeu was also recognized as InN_MGE and InN_CGE based on their origins, respectively (Figures 5B and 5C). The cell proportions of each cluster were comparable between WT and lissencephaly brains. However, we detected a significant deviation in cell proportion between MU and MD in lissencephaly brains, indicating that the cellular composition was markedly altered in the upper band and the deep band in *DCX^-/Y^* (Figures 5C and S3D).

**Figure 5.**
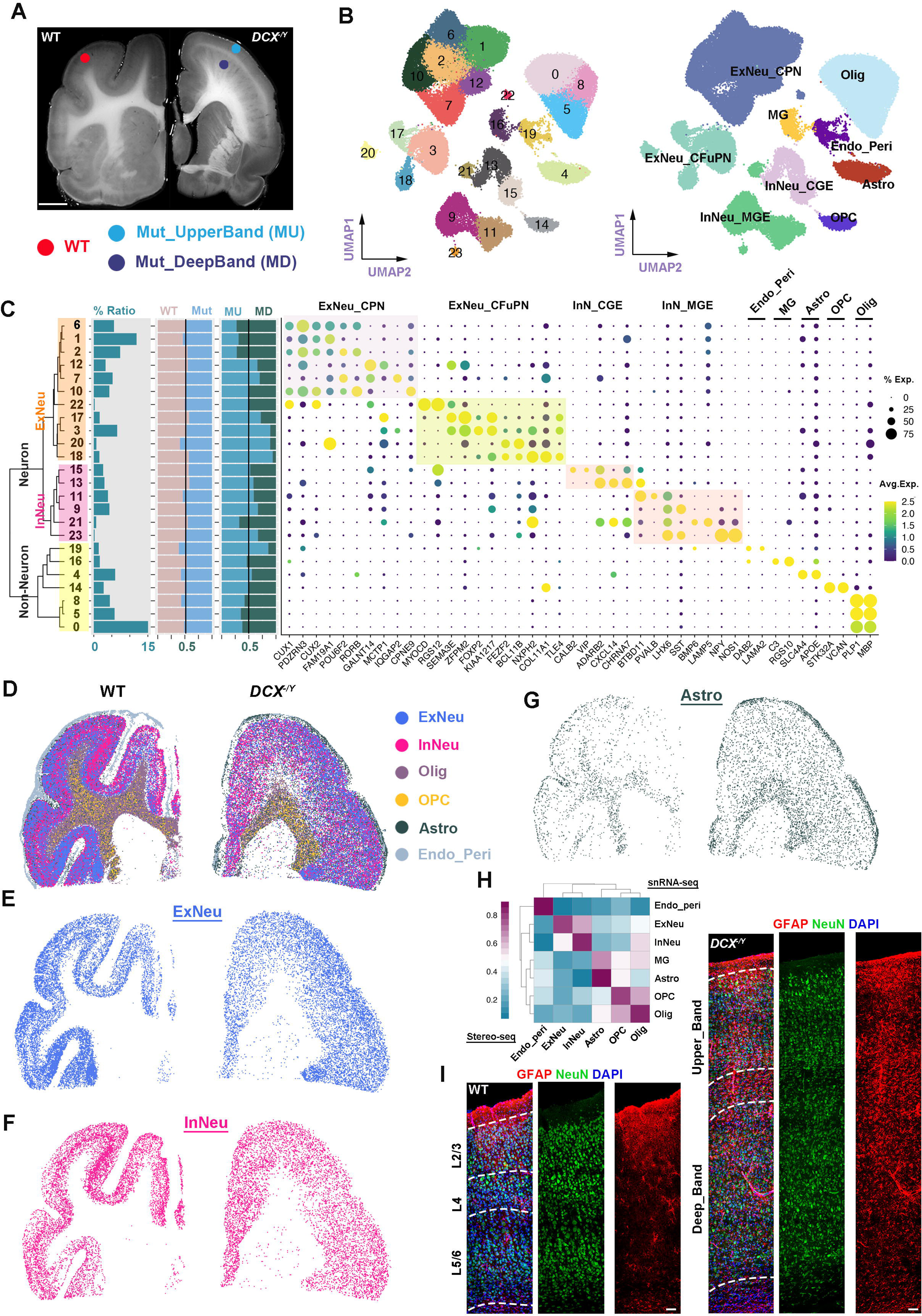
Single nuclei and spatial transcriptomic landscape of cell types in WT and *DCX* ferrets. (A) Sampling information used for snRNA-seq (scale bar, 500µm). (B) Clustering of individual cells visualized by UMAP (left). UMAP showing the annotation of cell types on the basis of gene expression (right). (C) Dot plot showing the average expression of marker genes for each cluster, and the phylogenic relationship of clusters is illustrated via dendrogram. Three bar plots showing the ratio of each cluster to total cells (left), WT to Mut (middle), and MU to MD samples (right). (D-G) Spatial transcriptome of major cell type in WT (left) and *DCX* knockout (right) ferret cortices at bin-80 resolution. (H) Correlation analysis of cell types between snRNA-seq and Stereo-seq. (I) The distribution of GFAP^+^ astrocytes in WT and *DCX* knockout ferrets (Scale bar, 100µm).

We also employed spatial transcriptomic analysis to investigate the spatial heterogeneity, with a mean gene number over 1900 in WT and 1200 in *DCX^-/Y^*at bin-80 resolution (Figure S3E). Initially, we observed distinct cortical and subcortical regions with their unique gene expression profiles (Figure S3F). We then only focused on cortical areas, including grey matter and white matter. In the cortices, we identified the major cell types and their locations (Figures 5D-5G and S3G). The cell types were consistent with those in the snRNA-seq dataset (Figure 5H). By comparing the distribution of each major cell type, we found that, besides excitatory and inhibitory neurons, astrocytes and oligodendrocytes also showed dramatically different patterns in WT and *DCX^-/Y^* (Figures 5D-5G and S3G). A dense band of oligodendrocytes was observed in the mutant, representing the white matter splitting the mutant cortex into MU and MD (Figure S3G). The distribution of astrocytes in *DCX^-/Y^*was altered compared to that in WT (Figure 5G). Immunofluorescence analysis showed that GFAP was uniformly and widely distributed throughout the entire cortices in *DCX^-/Y^*, but densely located in the outermost region of the cortices in WT, which confirmed the observation in the spatial transcriptomic dataset (Figures 5G and 5I). In summary, by using snRNA-seq and spatial transcriptome analysis, we reported the cellular landscape of the ferret neocortex. Our findings reveal significant alterations in the cellular composition and distribution patterns of major cell types, such as ExNeu, InNeu, and Astrocyte groups, between *DCX^-/Y^* and WT.

### Characteristics of Excitatory Neurons in lissencephaly

To find out the cellular and molecular differences between WT and *DCX* mutants, ExNeu cells were subset and grouped into 9 subtypes based on the DEGs (Figures 6A and S4A-S4C; Table S3). The 9 subtypes of excitatory neurons were identified in all samples; however, the ratio of each subtype varied significantly between MU and MD. Surprisingly, we found more deep-layer neurons (Layer5/6) were detected in MU, more upper-layer neurons (Layer2-4)were enriched in MD, indicating a disorganization of neuronal lamination in lissencephalic ferret brains (Figures 6A and S4A). Next, we investigated the localization of these layer-specific neurons using spatial transcriptomics. We found L23_ExNeu, L4_ExNeu, and L56_ExNeu located following regular lamination from the pial surface to the white matter in WT, but the lamination was completely disrupted in lissencephalic ferret brains, with more L23_ExNeu and L4_ExNeU in the deep part and more L56_ExNeu in the superficial part (Figure 6B), consistent with the cell ratio shown in snRNA-seq (Figure 6A). We validated the results observed in snRNA-seq and spatial transcriptome datasets by using antibodies against markers for upper- and deep-layer neurons in fluorescent immunostaining. A well-organized lamination was detected in WT, with CUX1 and SATB2 abundantly expressed in layer2-4, and CTIP2 and FOXP2 mostly expressed in layer 5 and layer 6 (Figure 6C). In *DCX^-/Y^* cortices, the localization of upper and deep-layer neuronal markers was disorganized. FOXP2 and CTIP2 positive cells were predominantly detected in the upper bands but were less abundant in the deep bands.

**Figure 6.**
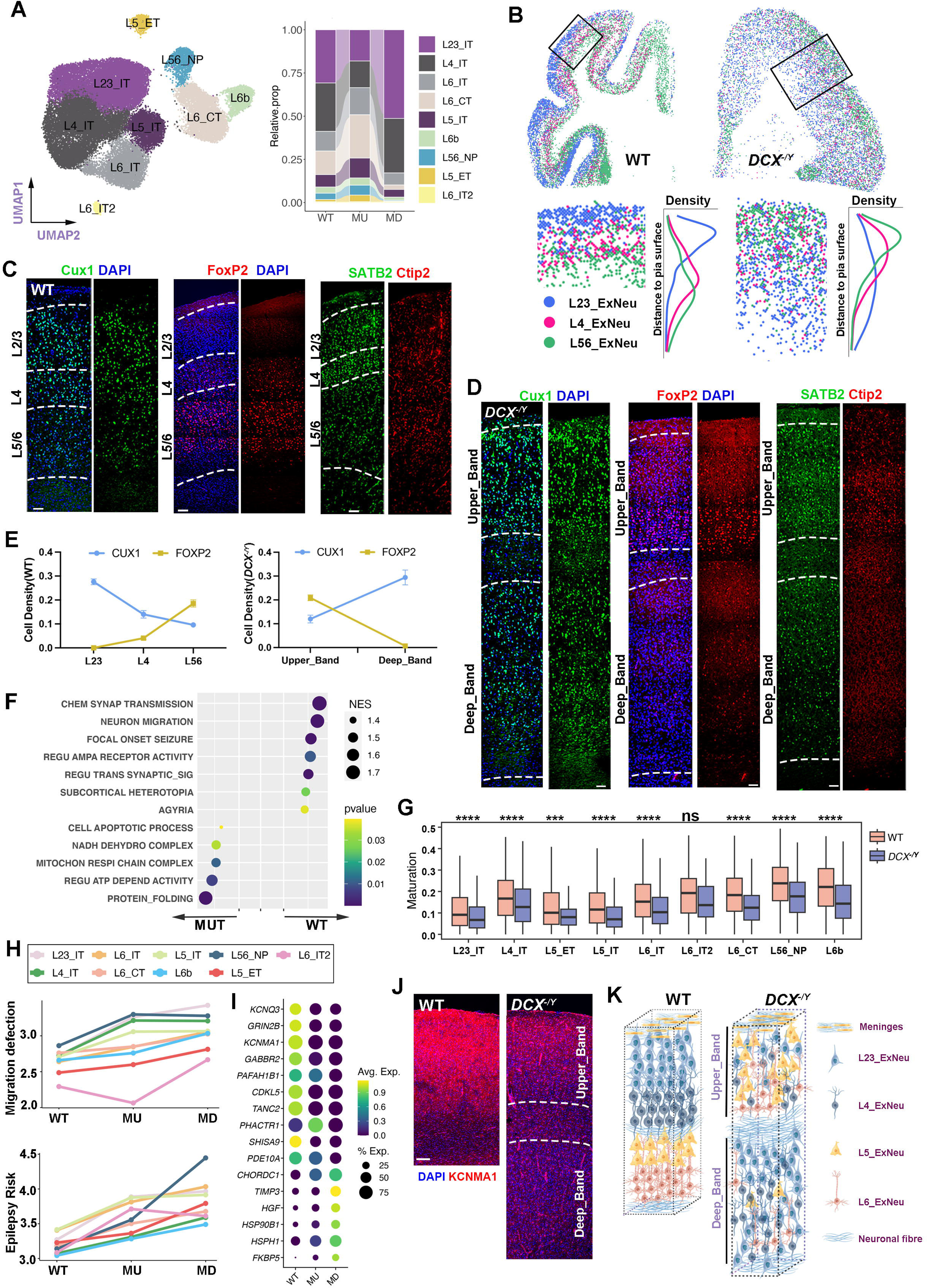
The lamination of excitatory neurons is disorganized in *DCX* knockout ferrets. (A) UMAP visualization of the ExNeu group. Cells are colored by the 9 annotated cell types (left) and the relative proportion of each cell type in WT, MU, and MD samples (right). (B) Spatial transcriptome showing L23_ExNeu, L4_ExNeu, and L56_ExNeu in the cortices of WT and *DCX* knockout animals (upper). Magnification images of the regions squared in the upper panels with the density of these three cell types (lower). (C and D) Immunofluorescent results showing the distribution of neuronal markers in WT and *DCX* knockout ferrets (Scale bars, 100µm). (E) Quantification results of the density of CUX1 and FOXP2 relative to DAPI in WT (n=4) and *DCX* (n=4). (F) Representative GSEA terms were significantly altered in WT and *DCX* knockout ferrets. (G) Boxplot showing the maturation scores of the 9 ExNeu subtypes in WT and *DCX* knockout samples calculated with CytoTRACE. The asterisk represents the p value of Kruskal-Wallis test in *DCX* compared to WT at maturation scores (**** for p < 0.0001, * for p <0.01). (H) Result of migration defection score (upper) and epilepsy risk score (lower) evaluated with AddModuleScore Function in Seurat for the 9 ExNeu subtypes in WT, MU, and MD samples. (I) Dot plot showing average expression of representative genes enriched in GSEA terms (H) in WT, MU, and MD samples. (J) Immunofluorescent result showing the expression level of KCNMA1, a seizure-risk gene, in WT, MU, and MD (Scale bars, 100µm). (K) Working model showing the disorganized lamination of excitatory neuronal subtypes in *DCX* knockout ferrets compared to that in WT (By Figdraw).

Conversely, both CUX1 and SATB2 positive cells were enriched in both the upper and deep bands (Figure 6D). Quantification analysis revealed a reversed distribution of marker genes in *DCX^-/Y^* compared to WT, as depicted in the spatial transcriptomic dataset (Figure 6E). In summary, we illustrated the cellular composition and the spatial distribution of excitatory neurons in WT and *DCX^-/Y^*, clearly indicating that the cytoarchitecture of mutant cortices was disrupted compared to that in WT (Figures 6A-6E).

We next applied Gene Set Enrichment Analysis (GSEA) to the DEGs to examine the functional alterations between WT and lissencephaly ferret brains (Figure S4D; Table S4). Notably, we observed that terms related to the well-known functions and phenotypes associated with DCX were down-regulated in *DCX^-/Y^*, such as agyria, subcortical heterotopia, and neuronal migration. Intractable epilepsy is a common symptom in human patients with *DCX* mutations and in mutant rodents.^29–32^The enriched terms also suggested that brain seizure (epilepsy), voltage-gated cation channel activity, and AMPA receptor activity were dysregulated (Figures 6F, S4E, and S4F), highlighting the molecular mechanisms underlying the onset of epilepsy.

Among the upregulated terms enriched in the mutant, chaperone-mediated protein folding and ATP-dependent activity implied the accumulation of a large amount of unfolded or misfolded polypeptides in *DCX* mutant neurons, which is observed in a variety of brain diseases.^33^ Mitochondrial respiratory chain related terms and the upregulation of the cell apoptosis process suggested the cellular stress may be high in *DCX* mutant cortices (Figures 6F and S4F).

The downregulated synapse-related signaling suggested neurons in *DCX* knockout animals might be less mature and/or less functional than those in WT. Thus, we further evaluated the maturation status with the Cytotrace package^34^ and found that nearly all cell types were less mature in both MU and MD compared to those in WT (Figures 6G and S4G).

Next, we asked whether there were gene expression differences between the upper band and deep band, and whether the neuron subtypes behaved differently compared to WT. To address this question, seizure-risk score and migration-defect score were calculated (Figure 6H). The results showed that both seizure-risk score and migration-defect score were increased in *DCX* knockout animals compared to those in WT across most neuronal subtypes, which indicated 1) the disorganized lamination in lissencephaly ferrets had a global impact on the gene expression of excitatory neurons but not on certain subtypes; 2) the cells in the MD were more severely affected than those in the MU (Figure 6H). The representative gene expression levels also verified that cells in MD were more severely affected (Figures 6I and S4H). The protein expression level of KCNMA1, a gene associated with an increased risk of seizures,^35^ was evaluated using a specific antibody in an immunofluorescent assay (Figure 6J). We found the expression level of KCNMA1 was high in the cortices of WT but dramatically decreased in *DCX* mutant cortices. In summary, we found the lamination of excitatory neurons was disrupted and partially reversed in *DCX* mutant cortices by using snRNA-seq, spatial transcriptome analysis, and immunofluorescent assays (Figure 6K). Analysis also provided molecular evidence and potential mechanisms for epilepsy in patients with *DCX* mutation.

### The distribution of interneuron couples with excitatory neuron

In the brain, a balance between excitatory and inhibitory neurons is pivotal to maintain homeostasis and ensure proper functionality. In *DCX^-/Y^*, we observed disruptions to the lamination and functionality of excitatory neurons (Figure 6). Here, we aimed to determine whether inhibitory neurons were also affected in *DCX^-/Y^* ferrets.

Initially, we further subdivided InNeu into 15 clusters (Figure 7A). The identities of these clusters were recognized based on their representative markers and DEGs: Martinotti and non-Martinotti cells (SST*^+^*), basket and chandelier cells (PVALB^+^), LAMP5 inhibitory neurons (LAMP5*^+^*), VIP inhibitory neurons (VIP^+^), Ivy cells (LAMP5^+^LHX6^+^), as well as long-range projecting neurons (SST^+^NPY^+^). We discovered that clusters 4 and 5 showed predominant expression of CXCL4, KIT, and CCK, but low LAMP5 expression, resembling L1 interneurons, including rosehip interneurons in human brains^36^ (Figures 7A, 7B, S5A, and S5B; Table S5).The relative proportion of each subtype revealed that the SST subtype was relatively enriched in MU, while the VIP subtype was abundant in MD (Figures 7C and S5A).

**Figure 7.**
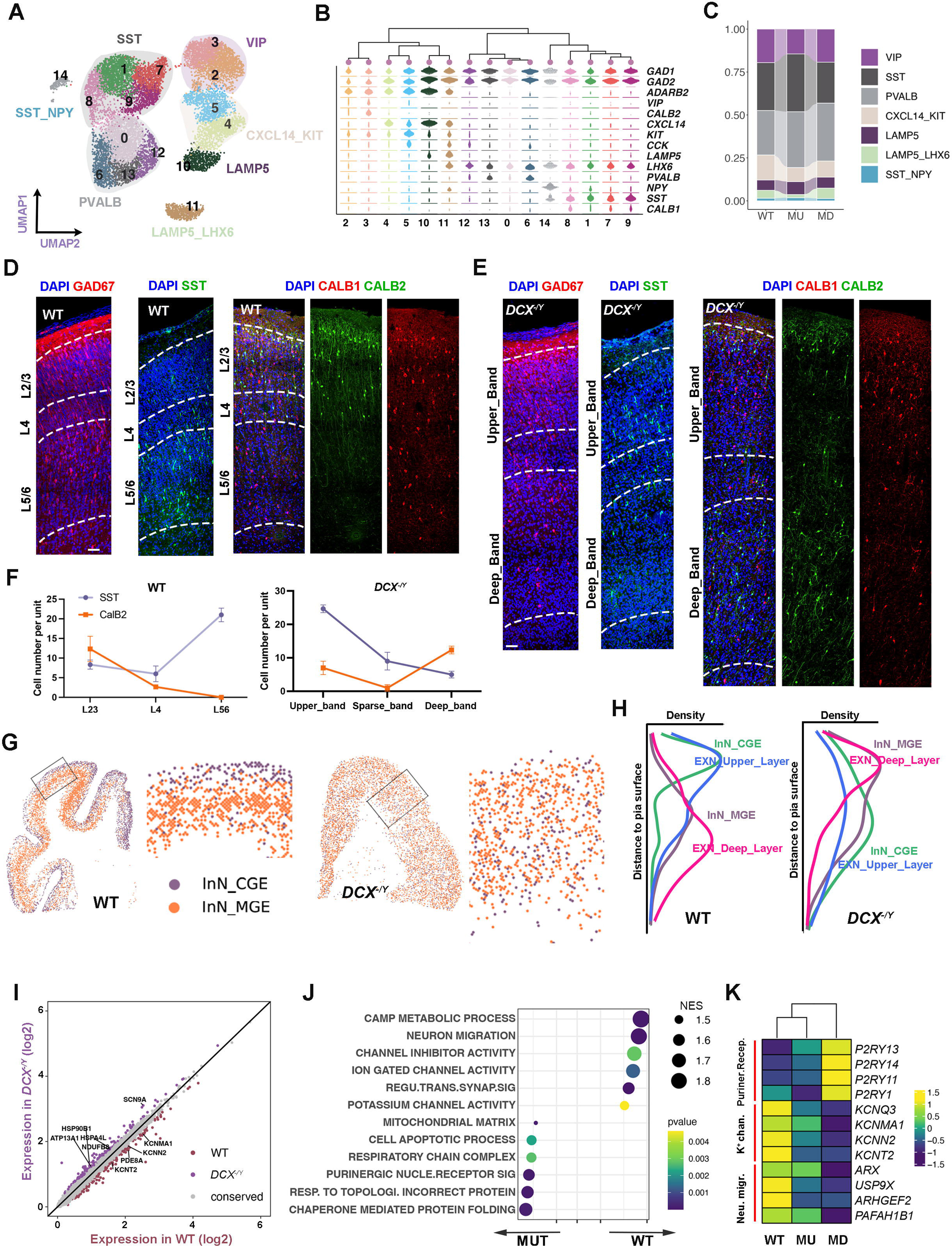
Interneuron profiles in *DCX* knockout ferrets. (A) Neuronal diversity of inhibitory neurons in WT and *DCX* knockout ferrets visualized via UMAP. Subtypes of inhibitory neurons were annotated based on the DEGs, which were highlighted with shadow colors. (B) Violin plots show the scaled expression of representative DEGs for each cluster, and a hierarchical tree is used to represent the relationship among the 15 clusters. (C) The relative proportion of InNeu cell types in WT, MU, and MD samples. (D and E) Immunofluorescent staining of adult ferret cortices for the interneuron markers GAD67, SST, CALB, and CALB2, showing disorganized distribution of interneurons in *DCX* knockout brains (Scale bars, 100µm). (F) Quantification analysis showing a similar pattern of inhibitory neurons with excitatory neurons in *DCX* knockout ferrets (n=4) compared to that in WT (n=4). (G) Spatial transcriptome showing InN_CGE, and InN_MGE in the cortices of WT and *DCX* knockout brains. Magnification images of the regions squared showing the distribution of the two cell groups. (H) Density plot showing the distance of the cell groups to the pial surface (G), including ExN_upper_layer, ExN_deep_layer, InN_CGE, and InN_MGE. (I) Scatter plot showing the differentially expressed genes in WT and *DCX* knockout ferrets. (J) Representative GSEA terms significantly changed in interneurons between WT and *DCX* knockout ferrets. (K) Heatmap showing average expression of representative genes enriched in GSEA terms (J) in WT, MU, and MD samples.

We then analyzed specific interneuron markers in the neocortex to evaluate their distribution. Disorganization in the distribution of SST, GAD67, CALB1, and CALB2 was observed in *DCX^-/Y^* compared to WT (Figures 7D-7F). Glutamate decarboxylase 67 (GAD67), encoded by the GAD1 gene, was almost evenly distributed across the cortex in WT, but predominantly in the upper band in *DCX^-/Y^* (Figures 7D and 7E). For specific interneuron subtypes in the WT, we observed SST positive cells primarily localized in the deep layer, in which deep-layer excitatory neurons were residing. Meanwhile, CALB2-positive cells were mainly found in the upper layer, where the upper-layer excitatory neurons were located. In contrast, within the *DCX* knockout brains, SST-positive cells were predominantly enriched in the upper band, a significant number of CALB2-positive cells were detected in the deep band of *DCX* knockout brains (Figures 7E and 7F).

Consistent with the immunofluorescence findings, our analysis of the spatial transcriptomic dataset showed a similar distribution pattern of InN derived from the medial ganglionic eminence (MGE) and the caudal ganglionic eminence (CGE) (InN_MGE and InN_CGE, respectively) in the WT. Specifically, InN_CGE was concentrated in the upper layer, while InN_MGE was predominantly located in the deep layer of the cortex (Figures 7G). The distribution pattern was disorganized in *DCX^-/Y^*. Interestingly, the distribution pattern of inhibitory neuron (InNeu) subtypes in WT and *DCX^-/Y^* showed a high accordance with the pattern of excitatory neuron (ExNeu) subtypes (Figures 6B-6E and 6K). This led us to speculate that the distribution of inhibitory neurons may be closely linked to specific excitatory neuronal subtypes.

To explore this possibility further, we applied cell density analysis to the spatial transcriptomic dataset. This allowed us to examine the spatial localization of the major cell types. We observed a close alignment of L23 and L4 ExN (ExN_Upper_layer) with InN_CGE inhibitory neurons, and L5 and L6 ExN (ExN_Deep_Layer) with InN_MGE in both WT and *DCX^-/Y^* (Figure 7H). Despite the disorganized distribution of ExNeu in *DCX^-/Y^*, these results suggest that specific inhibitory neuronal subtypes tend to colocalize with their corresponding excitatory partners.

We next conducted a GSEA on the DEGs to highlight the molecular changes in InNeu between WT and *DCX^-/Y^* globally and subtype specifically (Figures 7I and S5C; Table S6). Given the limited population of the SST^+^NPY^+^ subtype, we focused our GSEA term enrichment on the remaining subtypes (Figure S5D). Our findings revealed dysregulation of genes related to neuron migration, ion channels, synapse-related pathways, as well as mitochondrial functionality, and protein folding in *DCX^-/Y^* interneurons. More specifically, we detected a decrease in CAMP metabolism processes and potassium channel activity, along with an increase in the G protein-coupled purinergic nucleotide receptor pathway in *DCX^-/Y^*(Figures 7J and S5D), the dysregulations of these pathways were reportedly involved in the onset of brain seizure.^37,38^ The representative gene expression and gene-set scores underline our findings that cells in the deep band were more severely impaired than those in the upper band in *DCX^-/Y^*(Figures 7K and S5E), as the result observed in excitatory neurons (Figures 6H and 6I).

In conclusion, we have presented a comprehensive cellular and molecular profiling of InNeu in *DCX* knockout ferrets. Interestingly, we observed disorganization in the spatial distribution of interneuron subtypes. Furthermore, we found that the localization of interneurons was coupled with excitatory neurons in a cell type-specific manner. More specifically, interneurons derived from the MGE were predominantly coupled with layer 5/6 excitatory neurons, while interneurons derived from the CGE were mainly associated with layer 2-3 excitatory neurons.

## Discussion

In this study, we have systematically examined the phenotypes, cytoarchitecture, and molecular signatures of *DCX* mutant ferrets’ cortex. To the best of our knowledge, this constitutes the first animal model that recapitulates the full spectrum of human patient syndromes from both morphological and pathological perspectives. Here, we also provide strong evidence that DCX plays roles not only in neurons but also in NPCs by modulating their proliferation and the proper extension of radial glial fibers. We also showed the radial distribution of interneurons in the neocortex linked to excitatory neurons in a cell-type-specific manner.

The role of DCX in regulating neural stem cell proliferation during corticogenesis is still somewhat enigmatic and contentious. Mutations of *Dcx* seemingly have a slight effect on neurogenesis in rodents due to its exclusive expression in neurons in mice. The influence is significantly pronounced in double mutants of *Dcx/Lis1* and *Dcx/Dclk.*^39,40^ The proliferation of neural progenitors in these mutants is decreased, a finding which contradicts results observed in patient-derived (*DCX* mutant) induced pluripotent stem cells (iPSCs).^11^ In the current study, we reported an increase in the proliferation of NPCs in *DCX* mutant cortices, providing supporting evidence for

*DCX’*s role in suppressing NPC proliferation. We also provided evidence that overexpression of Dcx in mice neocortices decreased the proliferation of neural progenitors. Additionally, *in vitro* studies have shown that *DCX* overexpression markedly reduces self-renewal in brain tumor stem cells derived from human primary glioma,^41,42^ as well as human embryonic stem cell-derived neuronal progenitors.^43^ These findings further underscore *DCX’*s function in inhibiting cellular proliferation. Besides in ferrets, DCX is also detected in the neural progenitors of the developing brains of primates (such as macaques and humans), which implied indispensable function of DCX in gyrencephalic animals.

Gyrogenesis, the formation of convolutions in the cerebral cortex, is a characteristic feature of the human brain, and the process behind the formation of gyri and sulci continues attracting scientific interest.^44^ The emergence of oRG, which generate more cortical neurons, is considered a key driver of brain folding. However, in our study, we observed an abundance of cortical neurons in *DCX* mutants, suggesting that a sheer increase in neuron numbers is insufficient for gyrification. The proper localization of neurons is another vital element for brain folding. At later stages of cortical neurogenesis, radial glial fibers assume a fan-shaped structure, providing a scaffold for neuronal migration and tangential dispersion.^45^ This stage coincides with the onset of folding structures on the brain surface. In our study, we observed that the basal processes of RG/oRG were truncated or disintegrated in *DCX* mutant ferrets, devastating the proper distribution and dispersion of cortical neurons. Less migration defect shown at P0 might be caused by the short distance from the pial surface to the VZ. However, when the distance increased rapidly from P0 to P7, truncated fibers and lamination disorganization were observed. This finding strongly suggests that DCX modulates radial glial fiber extension and stability to support neuronal migration and brain folding. Past studies have shown that while a single knockout of *Dcx* in mice has an undetectable effect on neuron migration in the mouse cortex, *Dclk* knockdown severely disturbs radial fiber organization. Double mutants of *Dcx* (expressed in postmitotic neurons) and *Dclk* (expressed in NPC) present a stronger phenotype than either single mutant.^40,46,47^ Taking into consideration the diverse phenotypes observed in *DCX* mutants among rodents, humans, and ferrets, we hypothesize that this discrepancy might be due to species-specific differences in DCX expression profiles.^21,47^

The proper migration and positioning of excitatory and inhibitory neurons are critical for maintaining brain functionality and microcircuitry. Inhibitory neurons are generated and undergo tangential migration from the ganglionic eminences, after which they radially position themselves in their final locations in the cortex. Broadly speaking, interneurons derived from the medial ganglionic eminence (MGE) tend to be more densely distributed in the deeper layers of the cortex, whereas interneurons derived from the caudal ganglionic eminence (CGE) preferentially inhabit the upper cortical layer. An increasing body of evidence has revealed the attractive and repulsive mechanisms by which interneurons migrate to the dorsal cortex.^48^ However, the precise distribution of interneurons in the cortex is still not fully understood.

Pioneering studies in *reeler* mice demonstrated that inhibitory neurons often pair with excitatory projection neurons that are generated at the same time.^49–51^ Arlotta’s lab reported that somatostatin (SST) and parvalbumin (PV) interneurons are recruited by corticofugal projection neurons (deep-layer neurons) in Fezf2 mice.^52^ Our findings in *DCX* ferrets not only support Arlotta’s viewpoint but also extend it by showing that CGE-derived interneurons are recruited by upper-layer excitatory projection neurons. However, the detailed mechanisms tuning the precise laminar localization of subtype-specific interneurons remain elusive. Future studies, such as utilizing single-cell RNA sequencing and single-cell mass spectrometry on subtypes of excitatory and inhibitory neurons at different developmental stages, may provide valuable molecular insights into the attractive and repulsive interactions that govern these processes.

### Limitations of the study

In this study, we reported disrupted expression of ZO-1 and altered mitotic cleavage planes in apical radial glial cells in DCX knockout animals. The mechanisms through which DCX modulates the expression and/or stability of ZO-1 are unknown. We have observed changes in gene expression and GSEA in *DCX* knockout animals. However, since DCX is not expressed in neurons during adulthood, the alterations are probably caused by the disorganized cortical cytoarchitecture rather than being directly affected by DCX. Further investigation is needed to understand the mechanisms.

## Acknowledgement

This work was supported by the National Key R&D Program of China (2020YFA0804000, 2019YFA0110101), the National Natural Science Foundation of China (NSFC) (32122037, 32192411 and 81891001), the Science and Technology Innovation 2030—Brain Science and Brain-inspired Intelligence Project of China (2021ZD0200102, 2021ZD0202403, 2022ZD0211900), CAS Project for Young Scientists in Basic Research (YSBR-013), the New Cornerstone Science Foundation and Changping Laboratory.

## Author contributions

X.W., W.W. and Q.W. conceived the project and designed the experiments. C.Y., B.W. L.S. and J.Z. prepared animal samples. C. Y. and W.W. performed immunostaining, imaging and quantification analysis. Z. L., S.Z. and X.Z. performed the snRNA-seq. S.W., W.W. and W.M. analyzed the snRNA-seq data. S. W. and B.Z. analyzed spatial transcriptome data. W.W, Q.W. and X. W. wrote the manuscript. All authors edited and proofread the manuscript.

## STAR METHODS

### KEY RESOURCE TABLE

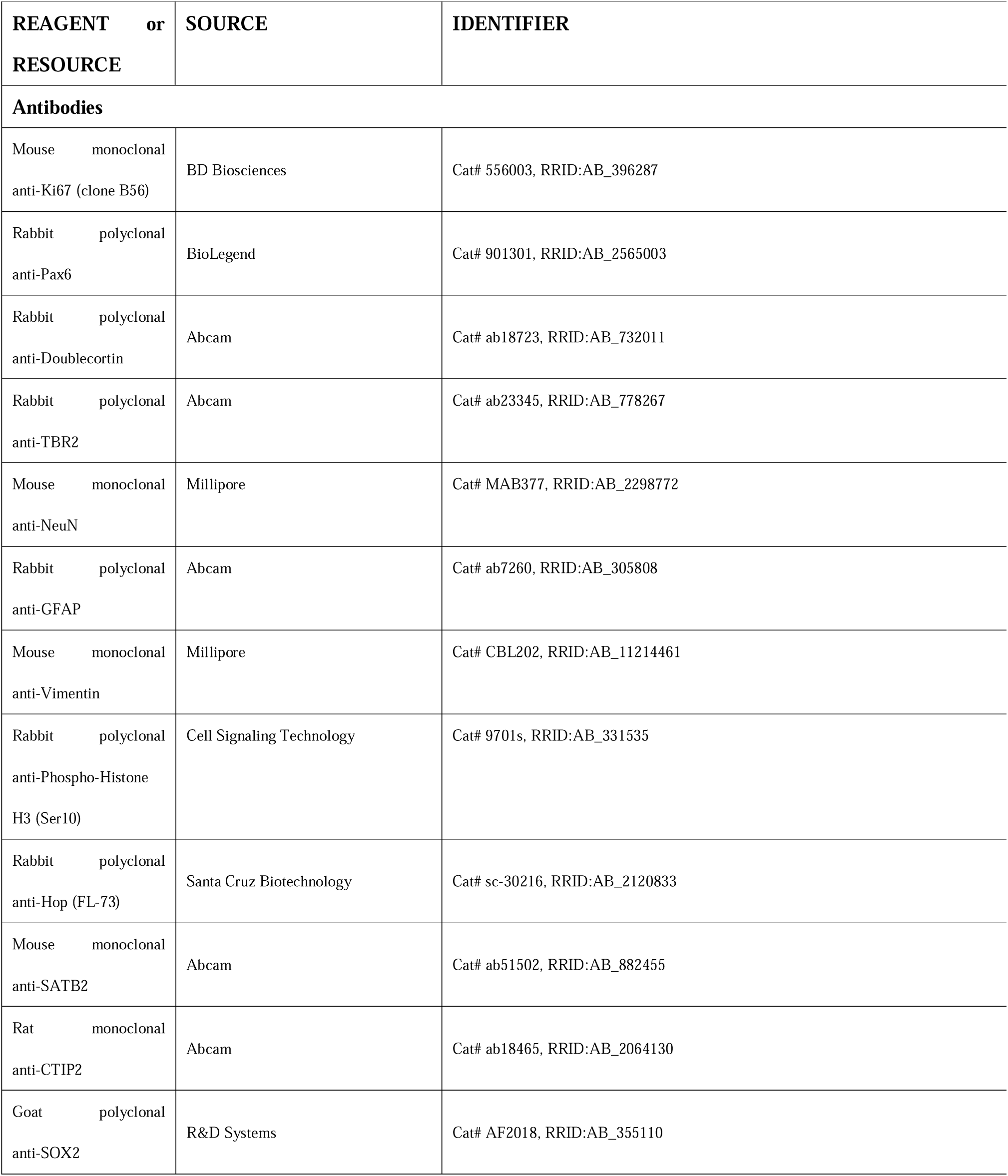
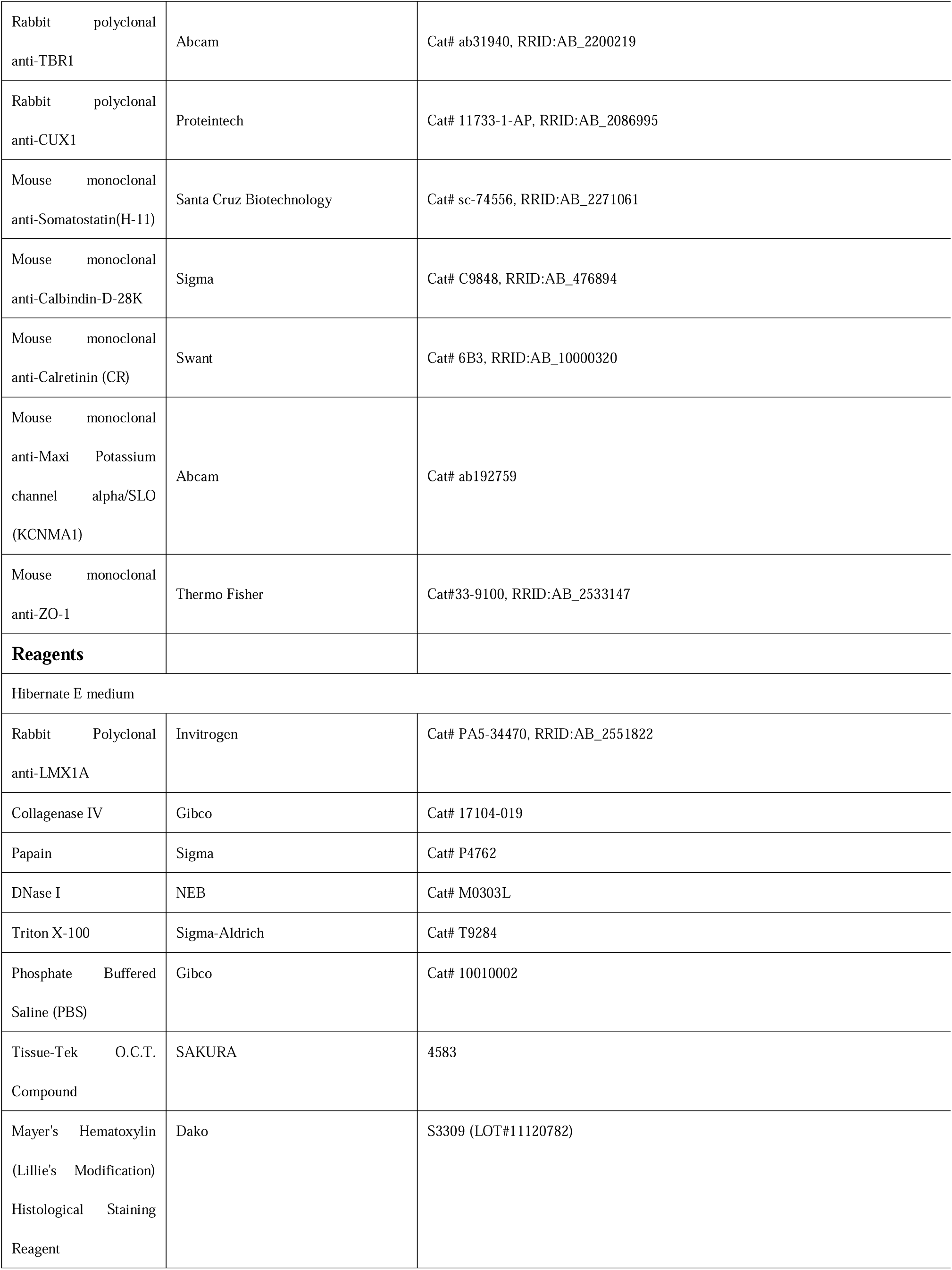
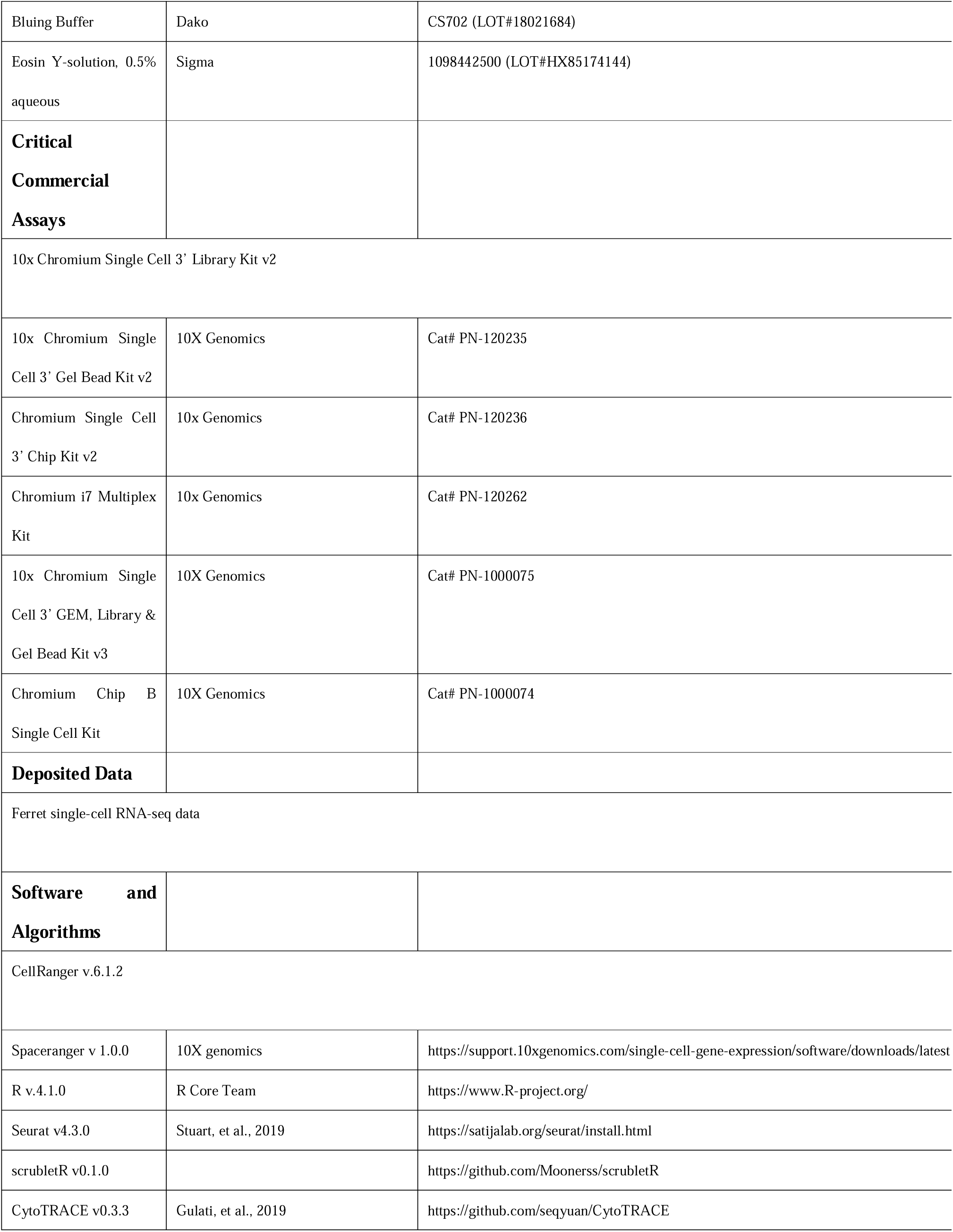
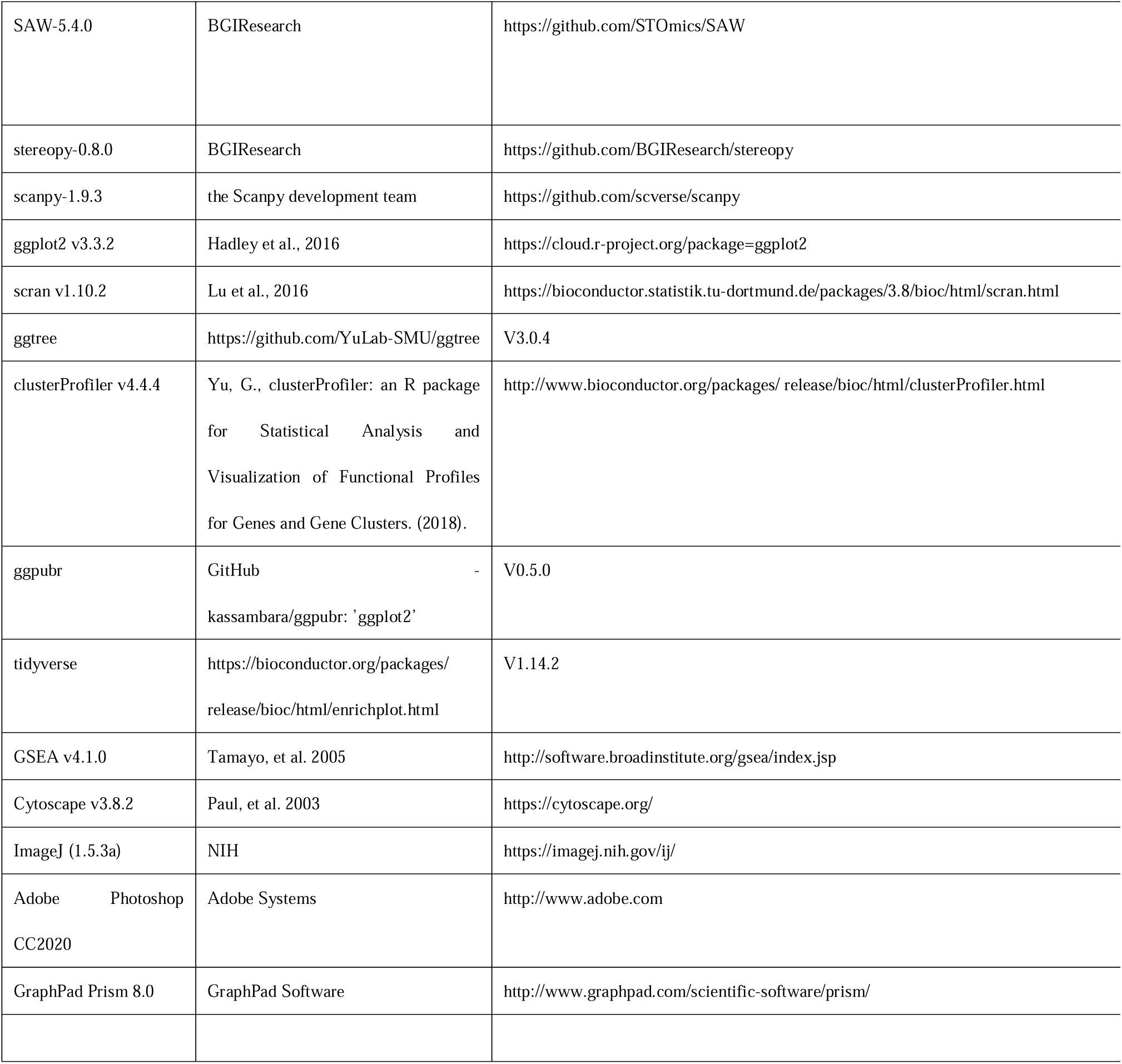

## RESOURCE AVAILABILITY

### Lead contact

Further information and requests for the resources and reagents may be directed to and will be fulfilled by the lead contact, Xiaoqun Wang.

### Materials availability

All materials used for stereo-seq and snRNA-seq are commercially available.

### Data and code availability

Sequencing data have been deposited in the Genome Sequence Archive (GSA) National Genomics Data Center (https://bigd.big.ac.cn/gsa-human) under accession number CRA011984 and the National Center for Biotechnology Information BioProjects Gene Expression Omnibus (GEO) under accession number GSE239781. There is no new code generated in this study. Further information and requests for data and code should be directed to and will be fulfilled by the corresponding author.

## EXPERIMENTAL MODEL AND STUDY PARTICIPANT DETAILS

### Animals

Crispr/CAS9-mediated knockout experiments were performed in Wuxi Sangosho Biotechnology CO, Wuxi, Jiangsu Province, China. Animal collection and experimental procedures followed national laws and international ethical and technical guidelines. The procedures were examined and approved in advance by the Committee on Animal Care of the Institute of Biophysics, Chinese Academy of Science. Detailed information for each genotype or age was provided in Table S1.

## METHOD DETAILS

### Ferret colony management and tissue collection

The *DCX* knockout ferret colonies and WT controls were bred and maintained at Wuxi Sangosho Biotechnology CO. All samples used in this study were collected at Wuxi Sangosho Biotechnology CO. and shipped to the Institute of Biophysics, Beijing, China.

Adult ferrets were anesthetized with over-dosed Zoletil^®^50 (Virbac) intramuscularly and perfused with cold artificial cortico-spinal fluid (ACSF). The whole brains were rapidly dissected on ice. The right and left hemispheres were separated and coronally divided into five parts that labeled as R1-R5 and L1-L5, respectively. The left parts were fixed in 4% paraformaldehyde (PFA) at 4 for 24h; the right hemispheres were fast frozen and stored in liquid nitrogen for snRNA-seq and stereo-seq experiments. For P0 and P7 samples, pups were deeply euthanized prior to brain removal. Brains were fixed in 4% PFA overnight for further use.

### *In utero* electroporation

The Dcx coding sequence of mice was cloned into pCAG-GFP plasmid (pCAG-DCX-GFP). The construct was verified by sequencing. DNA solutions were mixed in 10 mM Tris, pH 8.0, with 0.01% Fast Green. *In utero* microinjection and electroporation was performed at E13.5, using timed pregnant females. Forceps-type electrodes (BTX, ECM830) with 9 mm pads were used for electroporation (five 50-ms pulses of 30 V). Fetuses were examined at E16.5 for further analysis.

### MRI

All data were acquired in a SIEMENS Prisma plus Xcelerate 3.0 T MRI system. Anatomical images were acquired with the following T2 Rapid Acquisition with Relaxation Enhancement (RARE) sequence: repetition time (TR) =6580 ms, echo time (TE) = 85 ms, number of averages = 10, magnetic field strength = 3, flip angle = 150°, acquisition matrix = 320 x 190, matrix size = 256 x 128, FOV = 2.9 × 5.0 cm2, slice thickness = 1.5 mm, spacing between slices = 1.65mm.

### Nissl Staining

Samples were washed in PBS to remove OCT and immerse slices in Cresyl Violet 40 min-1 h 37, washed in 50% ethanol for 1min, 70% ethanol for 2 min and 100% ethanol for 30 s. Finally, samples were washed in xylene for 2x5min, and mounted with Fluoromount-G (Southern Biotech).

### DiI labeling

Embryo and adult ferret slices were fixed in 4% paraformaldehyde (PFA). DiI crystal (1,1’-dioctadecyl-3,3,3’,3’-tetramethylin-docarbocyanine-perchlorate; D-282, Molecular Probes) was applied to the ventricular zone or pial surface, which allowed it to diffuse throughout the tissues in PFA for at least 30 days (adult) at 37 .

### Fluorescent immunohistochemistry

The immunofluorescent assay was performed as previously described. Briefly, PFA-fixed ferret brain slabs were infiltrated in gradient sucrose solutions in PBS, and embedded in optimal cutting temperature (OCT) compound. Brains were sectioned at 10 µm thickness on a Leica Cryostat and stored at -80.

The slides were returned to room temperature and washed in 0.1M PBS for 30 mins, followed by antigen retrieval in sodium citrate (5uM, PH 6.0) at 90 for 10 min. Subsequently, samples were incubated overnight at 4 °C with 5% normal donkey serum (NDS) containing the primary antibodies (Star protocols). The secondary antibody incubation and DAPI nuclear staining were performed at room temperature for at least 2 h. Finally, samples were washed again in 0.1% PBST for 1h and mounted with Fluoromount-G (Southern Biotech). Images were obtained with Olympus FV3000.

### Brain sample dissociation and nuclei preparation

Somatosensory cortices were dissected carefully from fresh-frozen tissues stored in liquid nitrogen. Samples were minced into pieces <5 mm and then homogenized using a glass dounce tissue grinder (Sigma, D8938) in 2 mL of Nuclei EZ lysis buffer (Sigma, NUC-101) on ice. After two incubations on ice (with 4 mL of lysis buffer and 5 min each time), the homogenate was filtered through a 70 µm strainer, and then we used Debris Removal Solution (Miltenyi Biotech, 130-109-398) to perform density gradient centrifugation to clean the nuclear suspension according to the manufacturer’s protocol. Isolated nuclei were resuspended and washed with nuclei suspension buffer (NSB, consisting of 1× PBS, 0.1% BSA and 0.4 U/µL Ambion™ RNase inhibitor (Thermo Fisher, AM2684)) and filtered through a 35 µm cell strainer. Nuclei were counted using a hemocytometer and diluted to 1000 nuclei/μL for optimal 10× loading. Approximately 8000 nuclei were targeted and captured for each reaction.

### RNA library preparation for high-throughput sequencing

All the libraries were prepared on a 10X GENOMICS platform following the RNA library preparation protocols. Cells were partitioned into nanoliter-scale Gel Bead-In EMulsions (GEMs) using 10x GemCode Technology, where a common 10x Barcode labeled all the cDNA produced from the same cell. Primers containing an Illumina R1 sequence (read1 sequencing primer), a 16-bp 10x Barcode, a 10-bp randomer, and a poly-dT primer sequence were released and mixed with cell lysate and Master Mix upon dissolution of the single cell 30 gel bead in a GEM. The GEMs were incubated, and barcoded, full-length cDNA was generated from poly-adenylated mRNA by reverse transcription. Then the GEMs were broken, and the leftover biochemical reagents and primers were removed with silane magnetic beads. Before constructing the library, the cDNA amplicon size was optimized by enzymatic fragmentation and size selection. After that, P5, P7, a sample index, and R2 (read 2 primer sequence) were added to each selected cDNA during end repair and adaptor ligation. P5 and P7 primers were used in Illumina bridge amplification of the cDNA (http://10xgenomics.com). Finally, 150-bp paired-end reads were sequenced from each library using the Illumina HiSeq4000. Concerning the update of the Single Cell 3’ Regent Kits version, two libraries were prepared with Single Cell 3’ Regent Kits V3.

### Processing of scRNA-seq data from Chromium system

The sequence data were mapped to the ferret reference genome that we assembled and annotated ourselves to perform quality control and the read counting of genes using Cell Ranger (v.6.1.2) (http://10xgenomics.com) with default parameters. The gene-cell sparse matrix was generated for each sample by Cell Ranger software.

### Single-cell RNA-seq data processing

To detect potential doublets, the scrubletR (v0.1.0) (https://github.com/Moonerss/scrubletR) pipeline was performed on each sample by setting parameters ‘min_counts=2, min_cells=3, expected_doublet_rate=0.06, min_gene_variability_pctl=85, n_prin_comps=50, sim_doublet_ratio=2’. Cells with a computed doublet score greater than the doublet score threshold were identified as doublets and excluded from subsequent analysis. Next, the filtered cell-by-gene count matrix was loaded into Seurat for downstream analysis. We excluded cells that did not meet the following criteria: 1) cells with a number of expressed genes between 300 and 5000); 2) cells with a UMI (unique molecular identifier) between 200 and 15000); 3) cells with a percentage of mitochondrial counts smaller than 5. Genes that express in less than 5 cells were removed. Overall, 22366 genes and 77713 cells were retained for subsequent analysis. Next, we loaded the filtered count matrix into the CreateSeuratObject function to create a Seurat object, followed by log-normalization of the count matrix by the NormalizeData function. The top 2000 variable genes were identified using the FindVariableFeatures function. Then, principal component (PCA) analysis was performed using the RunPCA function. Batch effect correction was conducted on the principal components with function Harmony. Unsupervised clustering was performed with the functions FindNeighbors and FindClusters. Uniform manifold approximation and projection (UMAP) was employed for visualization of clustering with the RunUMAP function.

### DEG identification and cell type annotation

Differential gene expression analysis among clusters was performed using the Seurat FindAllMarkers function. Genes with adjusted P-values <0.05 were defined as differentially expressed genes (DEGs). Using respective classic markers, we identified 35 clusters including 7 major cell types: exciting neuron (ExN), inhibitory neuron (InN), neurons, oligodendrocyte precursor cells (OPC), oligodendrocyte (Olig), microglial cells (MG), astrocytes (AST) and endothelial cells/pericytes (Endo_Peri).

### Enrichment analysis of function pathways

To identify critical genes or function pathways between WT and *DCX* knockout ferrets in Neuron, gene set enrichment analysis (clusterProfiler v4.4.4) was performed to identify gene sets with statistically significant differences. Then the gene sets were tested in the Molecular Signature Database (MSigDB) C5 (ontology gene sets) that collection is divided into two subcollections, the first derived from the Gene Ontology resource (GO) which contains biological Process (BP), cellular Component (CC), and molecular Function (MF) components, and a second derived from the Human Phenotype Ontology (HPO). Gene sets enriched in several pathways were identified, including neuron migration, focal onset seizure, subcortical heterotopia and cell apoptosis. For each group comparison, the average gene expression was smoothed across cell types for the gene sets from each signaling pathway, as well as for the single representative gene. To observe the difference of pathway between WT, MU, and MD neurons, a pathway score was performed using AddModuleScore function of Seurat.

### CytoTRACE analysis for neuron

We used CytoTRACE (version 0.3.3) analysis to predict maturation state of ExN subtypes (L23_IT, L4_IT, L6_IT, L6_CT, L5_IT, L6b, L56_NP, L5_ET and L6_IT2) in WT and DCX samples (MU and MD) with default parameter, then wilconx test was used to test for score differences between the two groups (Mut vs WT).

### Stereo-seq library preparation and sequencing

Tissue collection and processing. Tissues sampled from two ferret brain were embedded with pre-cooled Tissue-Tek OCT (Sakura, 4583) immediately and snap-frozen in prechilled isopentane using liquid nitrogen until the OCT was completely solid. Embedded tissues were transferred to a -80 freezer for long time storage. The embedded tissues were cut to a thickness of 10 µm using a Leika CM1950 cryostat and then placed either in glass slide for Nissl staining or pre-chilled Stereo-seq capture chips for Stereo-seq procedures. Stereo-seq experiments were performed as previously described. Firstly, Stereo-seq capture chips were washed with NF-H2O supplemented with 0.05 U/µL RNase inhibitor (NEB, M0314L) and dried at room temperature. Then cryosections were adhered to the surface of the Stereo-seq capture chips and incubated at 37 for 5 min. Chips with sections were fixed in pre-cooled methanol at -20 for 40 min.

In situ reverse transcription. After fixation, chips with tissues were taken out and dry in the air. Then chips were washed with wash buffer (0.1 × SSC buffer [Thermo, AM9770] supplemented with 0.05 U/μl RNase inhibitor [NEB, M0314L]), tissue sections placed on the chips were permeabilized using 0.1% pepsin (Sigma, P7000) in 0.01 M HCl buffer, incubated at 37 °C for 6 minutes. Permeabilization reagent was then removed, and chips were washed with wash buffer. RNA released from the permeabilized tissues and captured by the DNB was reverse transcribed at 42 °C for 2 h using SuperScript II (Invitrogen, 18064-014, 10 U/μl reverse transcriptase, 1 mM dNTPs, 1 M betaine solution PCR reagent, 7.5 mM MgCl2, 5 mM DTT, 2 U/μl RNase inhibitor, 2.5 μM Stereo-seq-TSO [5-CTGCTGACGTACTGAGAGGC/rG//rG//iXNA_G/−3]) and 1 × First-Strand buffer. After that, tissues were washed twice with wash buffer and removed from the chips with tissue removal buffer (10 mM Tris-HCl, 25 mM EDTA, 100 mM NaCl, and 0.5% SDS) at 37 °C for 30 min. The chips were then treated with Exonuclease I (NEB, M0293L) at 37 °C for 1 h and washed twice with the wash buffer. The resulting first strand cDNAs on the chips were amplified using KAPA HiFi Hotstart Ready Mix (Roche, KK2602) with a 0.8 µM cDNA-PCR primer (5-CTGCTGACGTACTGAGAGGC-3), followed by incubation at 95 °C for 5 min, 15 cycles at 98 °C for 20 s, 58 °C for 20 s, and 72 °C for 3 min, and a final incubation at 72 °C for 5 min.

### Library construction and sequencing

The concentrations of the result cDNA products were quantified by the Qubit™ dsDNA Assay Kit (Thermo, Q32854) after purification using VAHTS DNA Clean Beads (Vazyme, N411-03, 0.6×). A total of 20 ng of products were fragmented using in-house Tn5 transposase at 55 °C for 10 min, and then the reaction was stopped by the addition of 0.02% SDS. Fragmented products were amplified as described below: 25 μl of fragmentation product, 1 × KAPA HiFi Hotstart Ready Mix, 0.3 μM Stereo-seq-Library-F primer (/5phos/ CTGCTGACGTACTGAGAGG*C*A-3), and 0.3 μM Stereo-seq-Library-R primer (5-GAGACGTTCTCGACTCAGCAGA-3) in a total volume of 100 µl with the addition of nuclease-free H2O. The reaction was run as: 1 cycle at 95 °C for 5 minutes, 13 cycles at 98 °C for 20 seconds, 58 °C for 20 seconds and 72 °C for 30 seconds, and 1 cycle at 72 °C for 5 minutes. PCR products were then purified using VAHTS DNA Clean Beads (0.6× and 0.15×). Finally, the library was used for DNB generation and sequenced using MGI DNBSEQ-Tx sequencer at China National Gene Bank (CNGB) as followed: 35 bp for read 1 and 100 bp for read 2.

### Stereo-seq data processing

We use SAW pipeline (https://github.com/BGIResearch/SAW) to analyze the raw data of Stereo-seq. Briefly, the identity of coordinates was mapped the designed coordinates matrix of the in situ captured chip which allowing 1 base mismatch and UMIs with N base or more than 2 bases with quality lower than 10 were filtered out. The retained reads were then aligned to the ferret genome using STAR, Mapped reads with MAPQ >= 10 were counted and annotated to their corresponding genes using the handleBam and used to generate a gene expression profile matrix. We divided the gene expression profile matrix into non-overlapping bins covering an area of 50 × 50 DNB, and the transcripts of the same gene aggregated within each bin. After this step, data were generated into GEF format for analysis by Stereopy (https://github.com/STOmics/Stereopy).

In the process of Stereopy analysis, we use the read_gef function to scan the GEF file setting bin=80 and use the filter_cells function to filter the bins (min_gene=100, min_n_genes_by_counts=3, max_n_genes_by_counts=2500). The Normalize_total, log1p and highly_variable_genes modules were separately applied for data standardization and high variable genes screening (min_mean=0.0125, max_mean=3, min_disp=0.5, n_top_genes=2000). PCA and UMAP dimensionality reduction analysis were performed as well. Finally, cluster was obtained by Leiden method. In order to observe the distribution of various types of neurons in space, we made tissue cuts according to coordinate points plotted in R-ggplot2.

### Statistics and reproducibility

All data obtained were repeated independently at least twice with similar results. Statistical analyses were performed by unpaired two-tailed Student’s t-tests and the multiple comparison test after ANOVA using GraphPad Prism software. Sample size and p values are given in the figure legends.

---

**Figure S1. The number and proliferation of NPCs are increased in *DCX* knockout ferrets, related to Figure 1**

(A) The sequences of two *DCX* knockout lines, Line1 is used in this study. Both lines introduce a stop codon in the second exon. Methionin, shown in the first place, is the 156^th^ amino acid.

(B) DCX shows abundant expression in the cortical plate (CP), the VZ, and the SVZ in WT, but no expression in knockout ferrets at P7. The expression of DCX is hardly detected in the VZ in mice at E14.5 and E16.5..

(C) Immunofluorescent images showing the expression pattern and colocalization of DCX with SOX2 and Vim at E35.

(D) The expression of DCX in the brains of developing monkeys and humans shows the colocalization with markers of RGs.

(E) Immunofluorescent staining results of pH3 show an increase of the number of proliferating cells in *DCX* knockout ferrets, compared to WT.

(F) The quantification of proliferating neural progenitors in the VZ, the iSVZ, and the oSVZ in WT and *DCX* knockout mutants at P0.

(G and H) More progenitors (Pax6^+^ in G) and proliferating progenitors (Ki67^+^Sox2^+^ in H) are detected in *DCX* knockout animals at P7. Scale bars:100µm.

**Figure S2. DCX modulates the extension of RG/oRG fibers and the migration of neurons, related to Figure 3**

(A) Nissl staining of WT and *DCX* knockout ferrets at P7. Arrow points out ectopic neurons which form a subcortical band.

(B and C) *DCX* knockout ferret brain surface is smooth, and the relative cortical length is shorter at P7, compared to WT (scale bar, 1mm).

(D) Abundant CUX1^+^ neurons are detected in *DCX* ferret, compared to WT at P7.

(E) VIM immunofluorescent staining shows no defect of RG/oRG basal fibers at P0.

(F) BLBP immunofluorescent staining shows no defect of RG/oRG basal fibers at P0.

(G) VIM immunofluorescent staining shows the defect of RG/oRG basal fibers at P7. Scale bars, 100µm.

**Figure S3. Transcriptome and spatial transcriptome features of somatosensory cortices in WT and *DCX ^-/Y^*, related to Figure 5**

(A and B) Quality control metrics for ferret samples; each dot represents a single cell. The number of genes (A) and counts (B) in each cell among various clusters in snRNA-seq.

(C) The expression of well-known marker genes for major cell types was visualized using UMAP.

(D) UMAP showing all clusters colored by major cell types in WT, MU, and MD samples.

(E) Quality control metrics for ferret samples; each dot represents a single cell. The number of counts (left) and genes (right) in each cell of various clusters in the stereo-seq section.

(F) The cell type annotation of the spatial transcriptomic section of the brain region in WT and *DCX* knockout samples.

(G) The cell annotation of the spatial transcriptomic section of major cell types in WT and *DCX* knockout samples.

**Figure S4. Transcriptional profiles of ExNeu in WT and *DCX* knockout ferrets, related to Figure 6**

(A) UMAP of snRNA-seq data showing all clusters colored by ExNeu subtypes in WT, MU, and MD samples.

(B) Dot plot showing the scaled expression of selected signature genes for ExNeu types; the size of the dots and the color bar were scaled with the average expression of the corresponding genes.

(C) Expression of marker genes for ExNeu types was visualized using UMAP.

(D) The DEGs of WT and *DCX* knockout ferrets, red dots representing the high profile in WT samples, blue dots representing the high profile in *DCX* knockout ferrets, and gray dots representing no significance.

(E) GSEA plot showing ‘Neuron migration’ and ‘Focal onset seizure’ were high in WT and low in *DCX* knockout ferrets.

(F) Representative GSEA terms changed in both MU and MD, compared to those in WT.

(G) Boxplot showing the maturation scores of the 9 ExNeu subtypes in WT, MU, and MD samples calculated with CytoTRACE. The asterisk represents the p value of Kruskal-Wallis test in *DCX* knockout ferrets compared to WT at maturation scores (**** for p < 0.0001, *for p <0.01).

(H) Correlation matrix of gene expression profiles in WT, MU, and MD.

**Figure S5. Transcriptional profiles of InNeu in WT and *DCX* knockout ferrets, related to Figure 7**

(A) UMAP visualization of InNeu subtype distribution in WT, MU, and MD samples.

(B) Expression of marker genes for InNeu subtypes visualized using UMAP.

(C) Volcano plot showing the different expressed genes of InNeu colored by subtypes in WT and *DCX* knockout ferrets, each dot representing a single gene.

(D) Representative GSEA terms significantly co-changed in subtypes of interneuron between WT and *DCX* knockout ferrets.

(E) Result of migration defection score, epilepsy risk score, and apoptosis score evaluated with AddModuleScore Function in Seurat for the subtypes of interneuron in WT, MU, and MD samples.

## Supplemental information

Figure S1-S6

Table S1. Summary of sample information for ferrets used in this study, related to all figures.

Table S2. Marker genes for the major cell type in ferret neocortex, related to Figure 3.

Table S3. List of marker genes highly expressed in ExNeu subtypes, related to Figure 4.

Table S4. List of differentially expressed genes in ExNeu across WT, MU, and MD, related to Figure 4.

Table S5. List of marker genes expressed in InNeu subtypes, related to Figure 5.

Table S6. Differentially expressed genes in InNeu between Mut and WT globally and cell-type specifically, related to Figure 5.

