## Supplementary figures and images for "*DCX* knockout ferret reveals a neurogenic mechanism in cortical development"

### SupplFigures_v13thJune.pdf

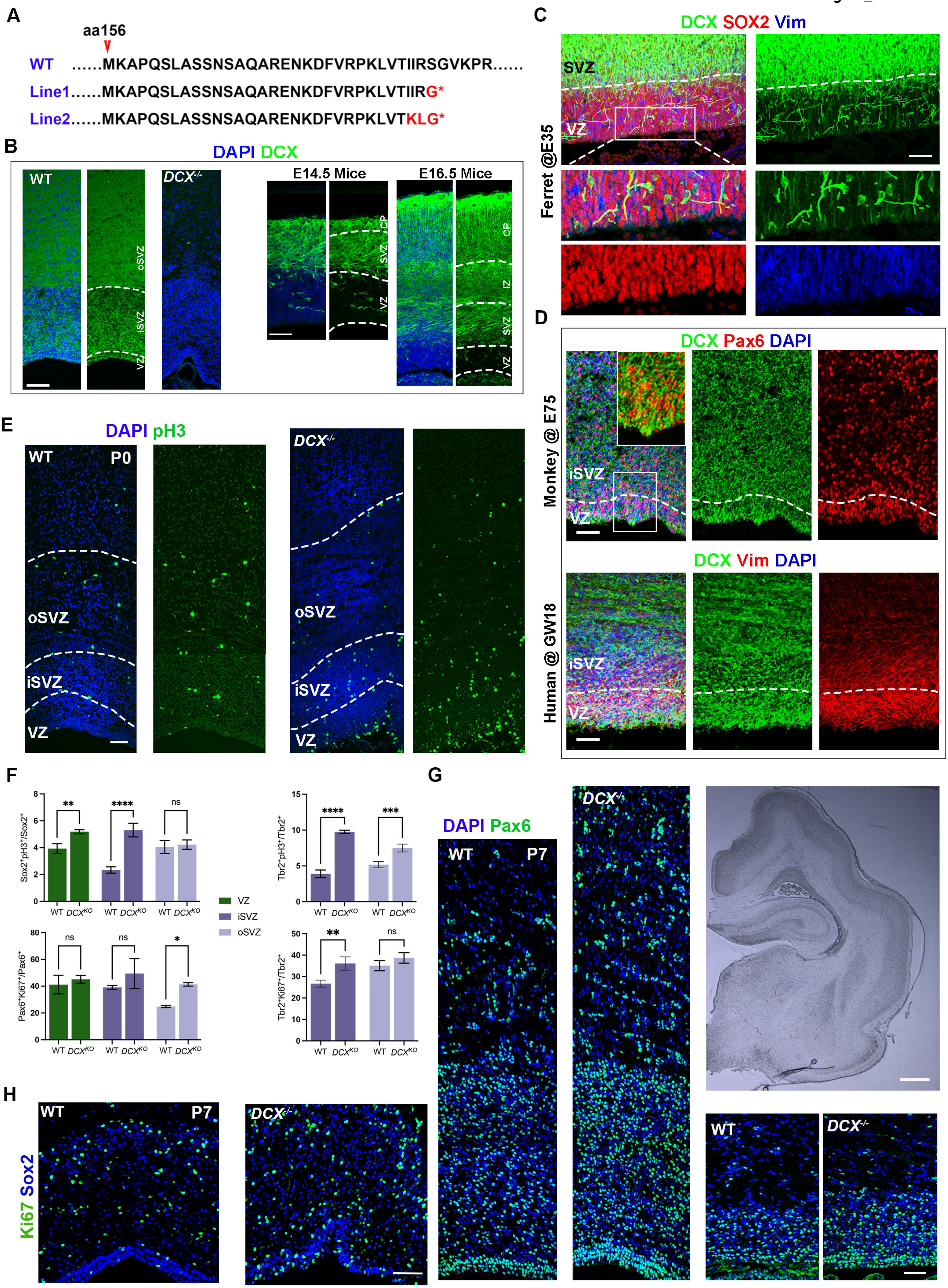

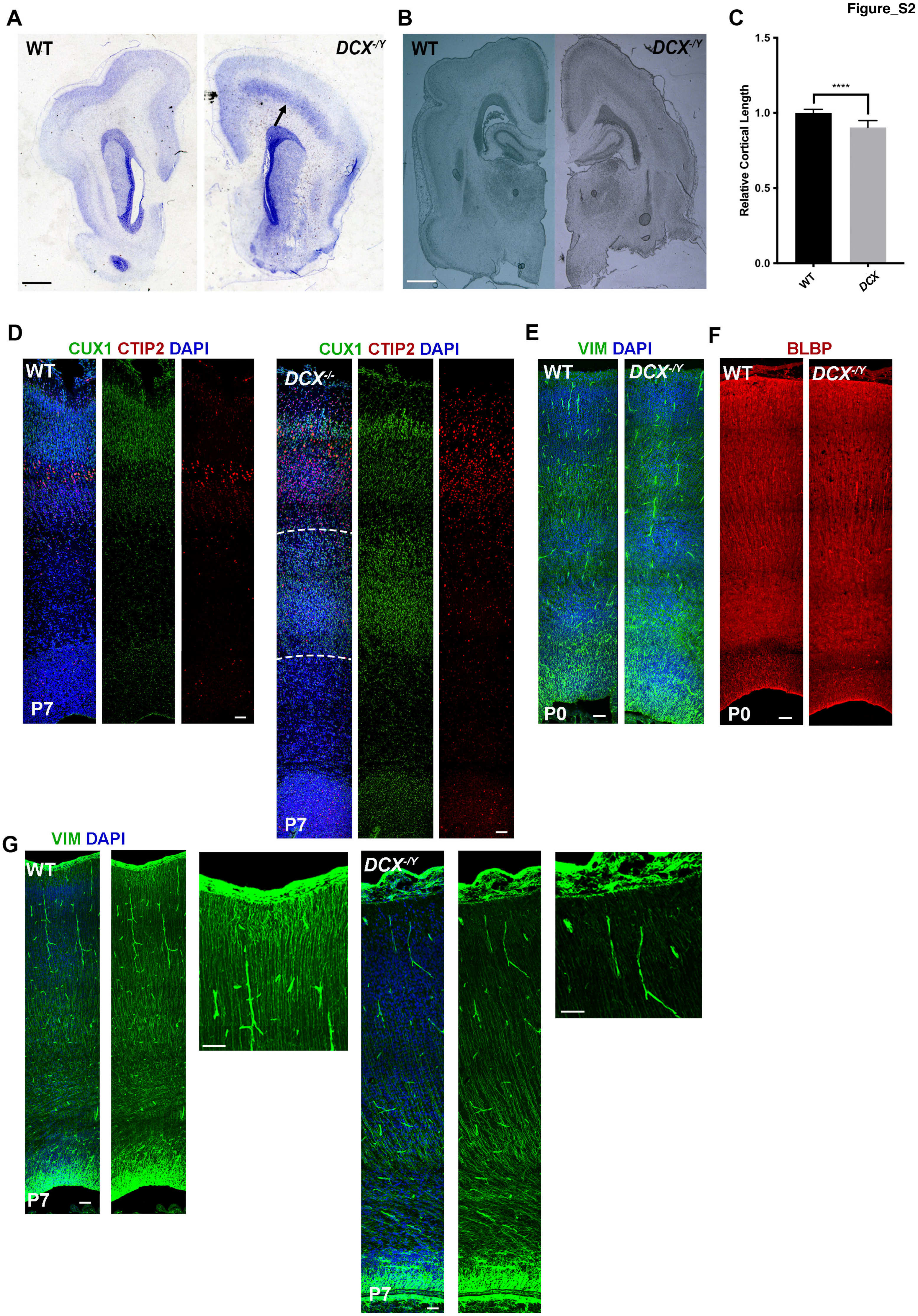

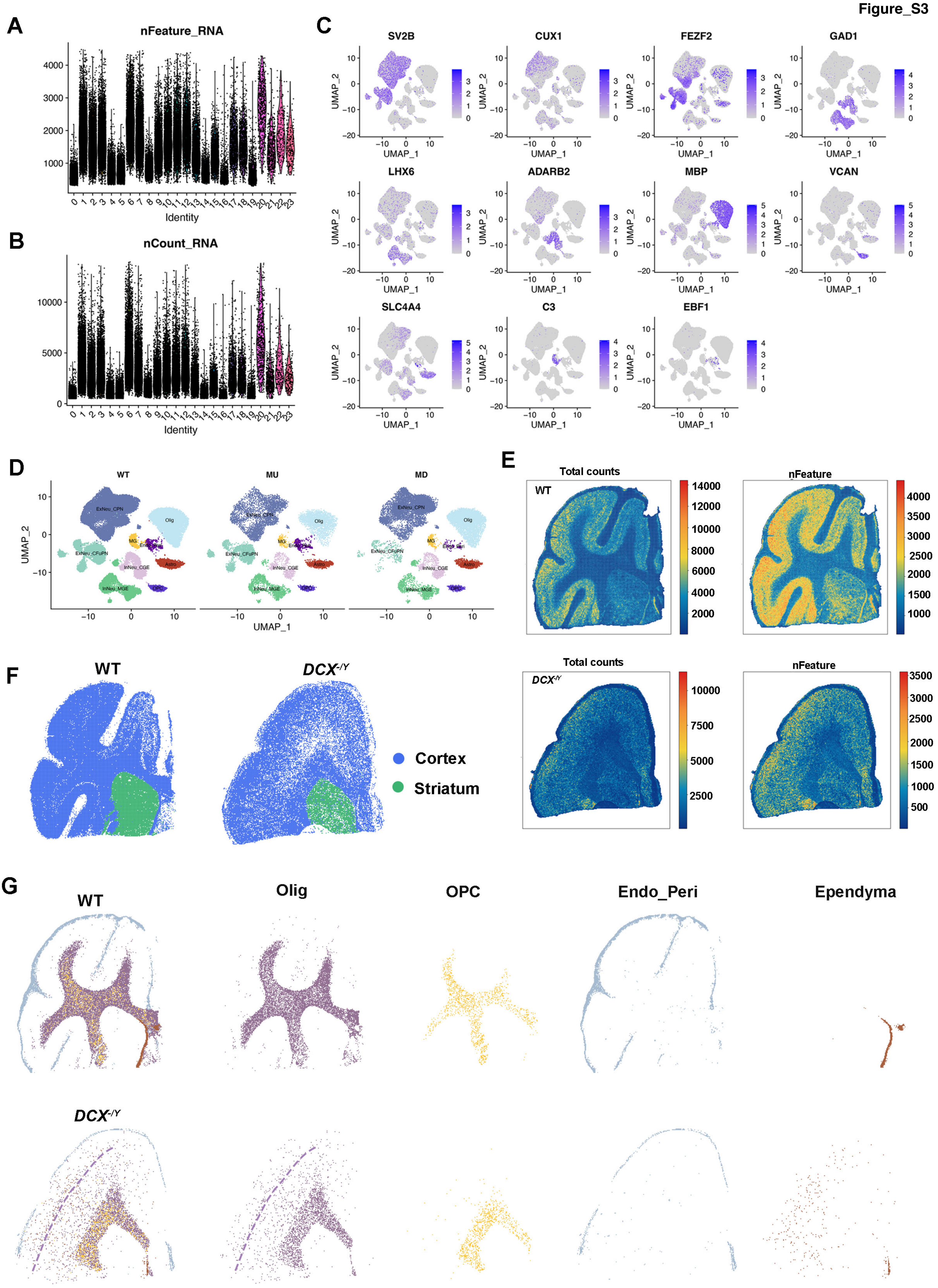

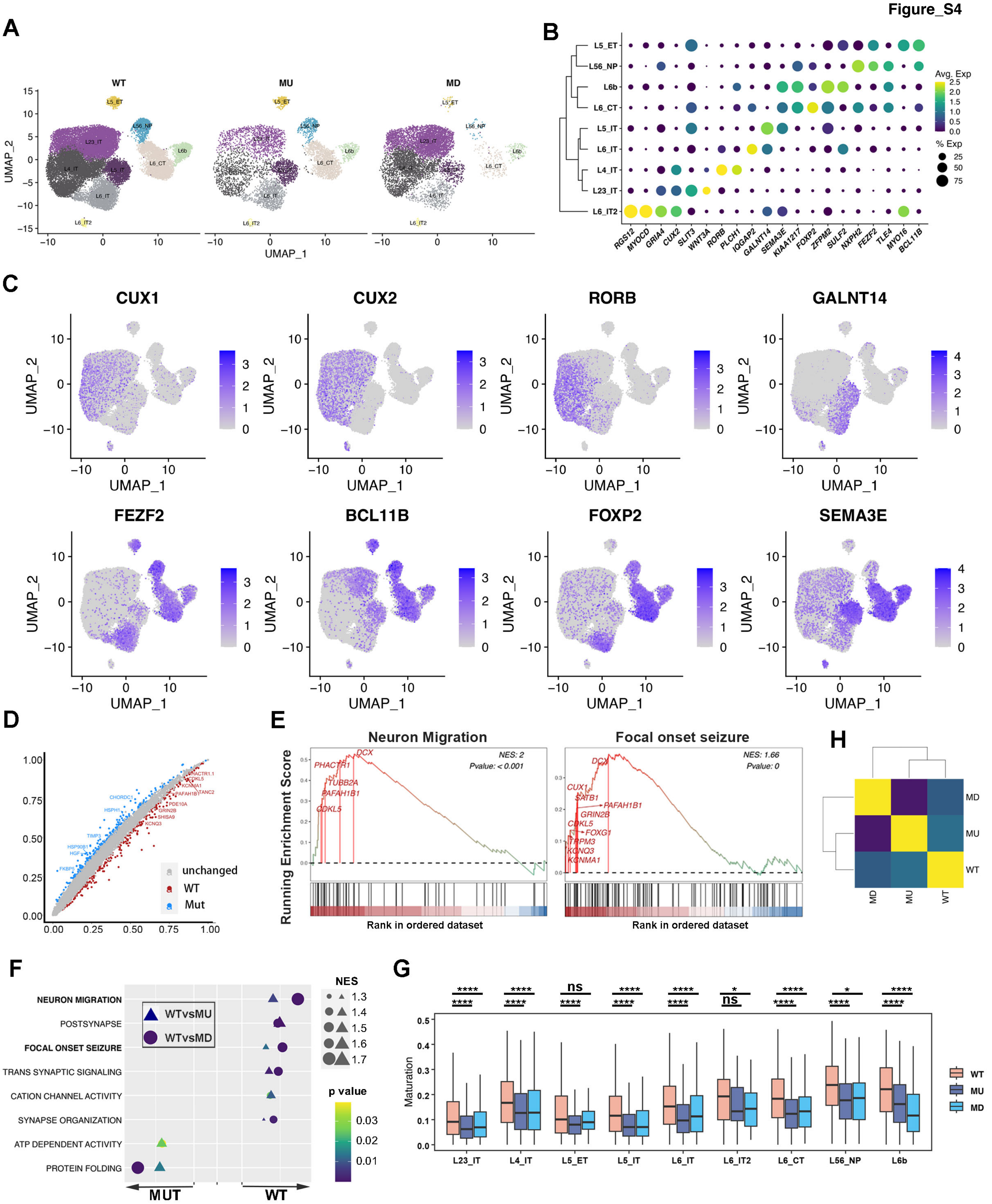

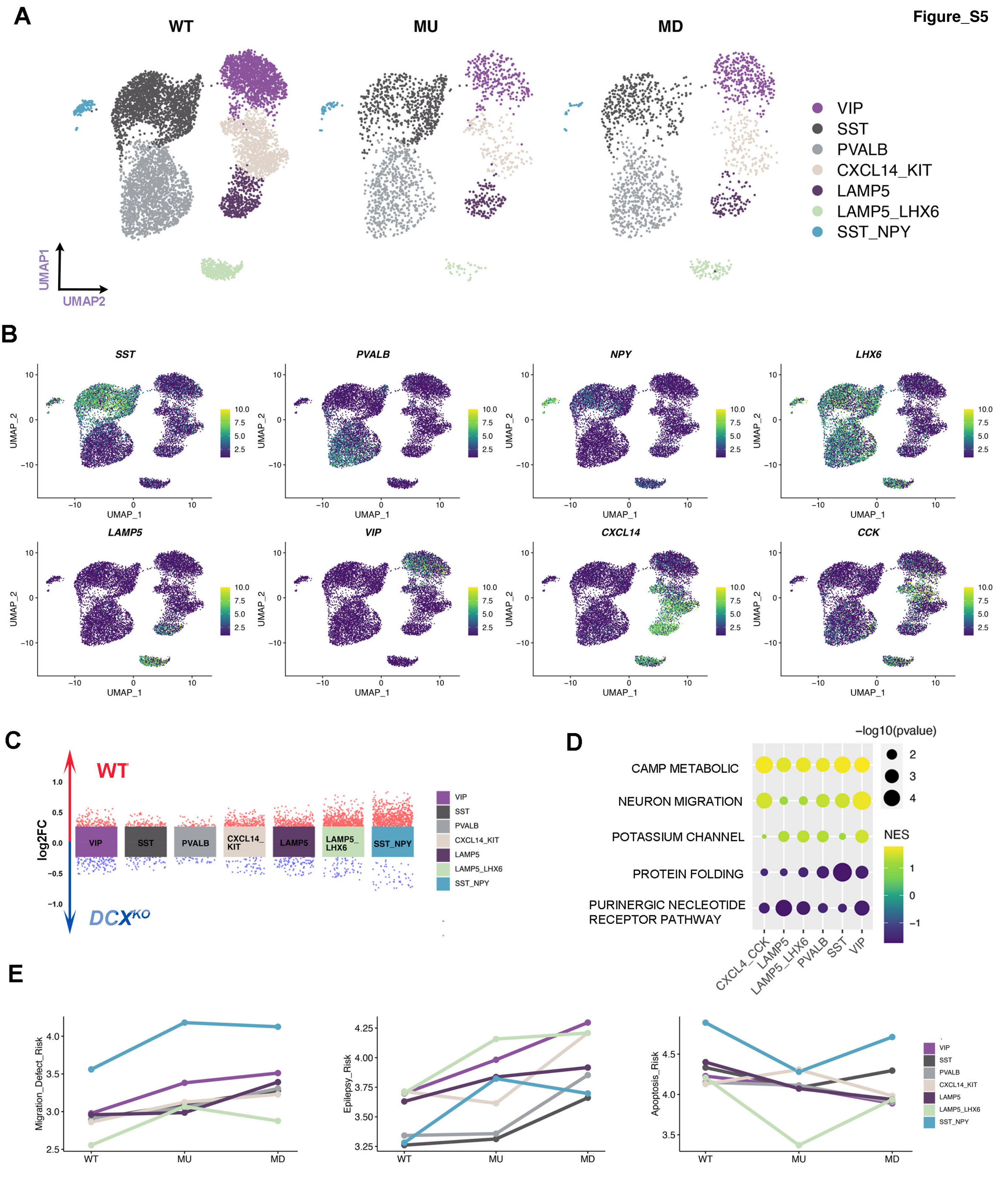
